# Within-host genomics of SARS-CoV-2

**DOI:** 10.1101/2020.05.28.118992

**Authors:** Katrina A. Lythgoe, Matthew Hall, Luca Ferretti, Mariateresa de Cesare, George MacIntyre-Cockett, Amy Trebes, Monique Andersson, Newton Otecko, Emma L. Wise, Nathan Moore, Jessica Lynch, Stephen Kidd, Nicholas Cortes, Matilde Mori, Rebecca Williams, Gabrielle Vernet, Anita Justice, Angie Green, Samuel M. Nicholls, M. Azim Ansari, Lucie Abeler-Dörner, Catrin E. Moore, Timothy E. A. Peto, David W. Eyre, Robert Shaw, Peter Simmonds, David Buck, John A. Todd, on behalf of OVSG Analysis Group, Thomas R. Connor, Ana da Silva Filipe, James Shepherd, Emma C. Thomson, The COVID-19 Genomics UK (COG-UK) consortium, David Bonsall, Christophe Fraser, Tanya Golubchik

**Author notes:** www.cogconsortium.uk. Full list of names and affiliations are in the Supplementary Material. Equal contribution (Katrina A. Lythgoe, Matthew Hall).

## Abstract

Extensive global sampling and whole genome sequencing of the pandemic virus SARS-CoV-2 have enabled researchers to characterise its spread, and to identify mutations that may increase transmission or enable the virus to escape therapies or vaccines. Two important components of viral spread are how frequently variants arise within individuals, and how likely they are to be transmitted. Here, we characterise the within-host diversity of SARS-CoV-2, and the extent to which genetic diversity is transmitted, by quantifying variant frequencies in 1390 clinical samples from the UK, many from individuals in known epidemiological clusters. We show that SARS-CoV-2 infections are characterised by low levels of within-host diversity across the entire viral genome, with evidence of strong evolutionary constraint in Spike, a key target of vaccines and antibody-based therapies. Although within-host variants can be observed in multiple individuals in the same phylogenetic or epidemiological cluster, highly infectious individuals with high viral load carry only a limited repertoire of viral diversity. Most viral variants are either lost, or occasionally fixed, at the point of transmission, consistent with a narrow transmission bottleneck. These results suggest potential vaccine-escape mutations are likely to be rare in infectious individuals. Nonetheless, we identified Spike variants present in multiple individuals that may affect receptor binding or neutralisation by antibodies. Since the fitness advantage of escape mutations in highly-vaccinated populations is likely to be substantial, resulting in rapid spread if and when they do emerge, these findings underline the need for continued vigilance and monitoring.

## Introduction

The ongoing evolution of SARS-CoV-2 has been the topic of considerable interest as the pandemic has unfolded. Clear lineage-defining single nucleotide polymorphisms (SNPs) have emerged (*1*), enabling tracking of viral spread (*2, 3*), but also raising concerns that new mutations may confer selective advantages on the virus, hampering efforts at control. Most prominently, there is increasing evidence that the D614G mutation (genome position 23403) in the Spike protein (S) increases viral transmissibility (*4, 5*) and N439K (genome position 22879) evades antibodies without loss of fitness (*6*). Most analyses have been focused on mutations observed in viral consensus genomes, which represent the dominant variants within infected individuals. Ultimately though, new mutations emerge within individuals, and hence knowledge of the full underlying within-host diversity of the virus at the population level, and how frequently this is transmitted, is important for understanding adaptation and patterns of spread.

The United Kingdom (UK) experienced one of the most severe first waves of infection, with over a thousand independent importation events contributing to substantial viral diversity during this period (*7*). In this study, we collected and analysed 1390 samples predominantly from symptomatic individuals (1173 unique individuals plus 93 anonymous samples) who tested positive for COVID-19 during the first wave of infection (March - June 2020; Table S1). The samples were collected by two geographically separate hospital trusts: Oxford University Hospitals and Basingstoke and North Hampshire Hospital, located 60 km apart. Using veSEQ, an RNA-Seq protocol based on a quantitative targeted enrichment strategy (*8*), which we previously validated for other viruses (*8*–*11*), we characterised the full spectrum of within-host diversity in SARS-CoV-2 and analysed it in the context of the consensus phylogeny.

We observed low levels of viral diversity within individuals, with evidence of strong within-host evolutionary constraint in Spike and other regions of the genome. Although within-host variants can be observed in multiple individuals in the same phylogenetic or epidemiological cluster, most viral variants are either lost, or occasionally fixed, at the point of transmission, with a narrow transmission bottleneck. These results suggest potential vaccine- or therapy-escape mutations are likely to rarely emerge or be transmitted from infectious individuals. Nonetheless, we identified Spike variants present in multiple individuals that may affect receptor binding or neutralisation by antibodies. Since the fitness advantage of escape mutations in highly-vaccinated populations is likely to be substantial, resulting in rapid spread if and when they do emerge, these findings underline the need for continued vigilance and monitoring.

### Detection of variants is influenced by viral load

Reliable estimation of variant frequencies requires quantitative sequencing, such that the number of reads is proportional to the amount of corresponding sequence in the sample of interest. The veSEQ protocol has been previously shown to be quantitative for a number of different pathogens (*9*), including acute respiratory viruses such as RSV (*10*). We demonstrated the same quantitative relationship holds for SARS-CoV-2. The number of uniquely mapped sequencing reads we obtained rose linearly with the number of RNA copies in serial dilutions of synthetic RNA controls (Fig. S1A, *r^2^*=0.87), and was consequently correlated with cycle threshold (Ct) values of clinical samples (Fig. S1B), indicating that veSEQ reads can be considered a representative sample of viral sequences within the input RNA. To calibrate our variant calling and minimise false discovery rates, we compared intrahost single-nucleotide variants (iSNVs) in re-sequenced controls with data for the stock RNA sequenced and provided by the manufacturer (Twist Bioscience) and masked sites vulnerable to *in vitro* generation of variants.

Next, we quantified the number of iSNVs in the full set of 1390 clinical samples at thresholds for identifying variants of between 2 and 5% minor allele frequency (MAF) (Fig. 1A). A minimum depth of at least 100 reads was also required to call an iSNV. For each threshold, we observed an inverse relationship between sample viral load (VL) and the number of detected iSNVs, but no association between mean MAF with number of mapped reads when no threshold was applied (*p*=0.291, linear regression, Fig. 1B). These observations can be partly explained by lower VL samples having fewer total observed reads, since the variance in observed MAFs is negatively correlated with read count (Fig. 1C). This is a straightforward probabilistic consequence of using repeated draws from a population to estimate the proportion of that population which has a discrete characteristic. However, this does not preclude the existence of biological mechanisms also contributing to greater intrahost diversity in low-VL samples, for example, if more variants are present later in infection when VLs are also lower. Since transmission appears to be more common at high VLs (*12*), variants observed in high VL samples are most likely to be available for transmission.

### Within-host variant frequencies are reproducible

Establishing reliable variant calling thresholds for clinical samples, where true variant frequencies are unknown, ideally requires re-sequencing of multiple samples from RNA to test for concordance. Working within the constraints of small volumes of remnant RNA from laboratory testing, we re-sequenced 65 samples, of which 27 replicate pairs generated sufficient read numbers (>50,000 unique mapped reads) for reliable minor variant detection. Intrahost single-nucleotide variants (iSNVs) with <2% MAF were generally indistinguishable from noise, whereas those >=3% MAF were highly concordant between replicates (Fig. 1D, Fig. S2).

Based on the above considerations, we identified a set of 583 iSNV sites that were observed (i) in high-VL samples with at least 50,000 unique mapped reads, (ii) at depth of at least 100 reads, (iii) with a MAF of at least 3%, and (iv) not observed to vary in synthetic RNA controls (Table S2; see Methods). Of these, we excluded the 18 sites which were variant in over 20 samples. Variants at these sites occurred at low frequency in many samples (Table S2), with some showing evidence of strand bias and/or low reproducibility between technical replicates (Fig. S2). Among the excluded sites was 11083, which was observed in 46 samples and is globally ubiquitous in GISAID data. From manual examination of mapped reads in our dataset, this appears to be due to a common mis-calling of a within-host polymorphic deletion upstream at site 11082, occurring in a poly-T homopolymeric stretch. If genuine, this homopolymer stutter may have a structural or regulatory role; however, methodological issues in resolving this difficult-to-map region cannot be ruled out. The remaining 565 sites were taken forward for variant analysis.

**Fig. 1.**
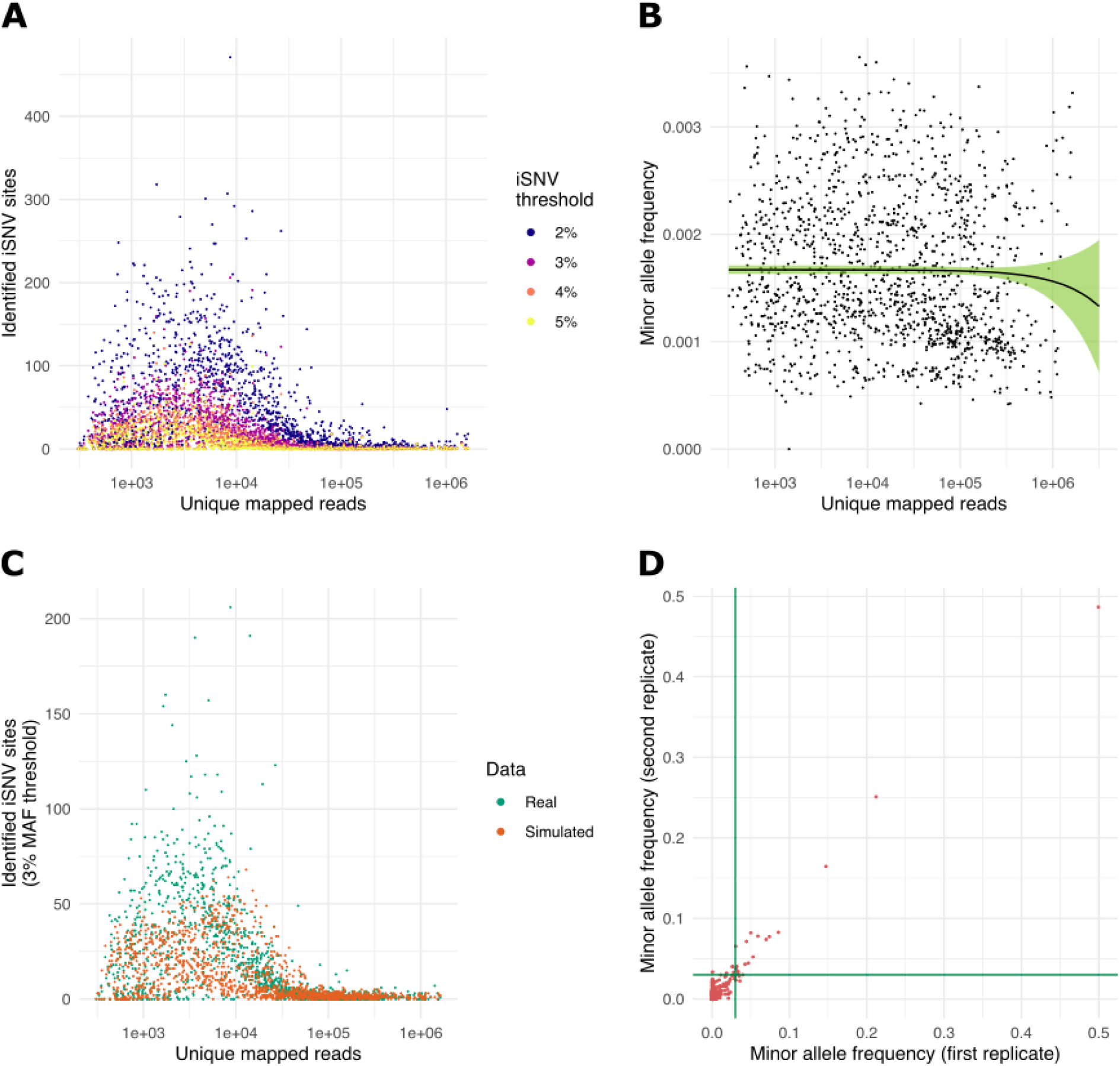
iSNV frequencies are reproducible. **A**: Distribution of number of identified iSNV sites at thresholds of 2-5% against number of unique mapped reads. **B**: Distribution of mean MAF against number of unique mapped reads. The black line is the estimated mean value by linear regression, with the green ribbon the 95% confidence interval. **C:** Distribution of number of identified iSNV sites at 3% from the real data (green) and from a simulation (orange). For the simulation ‘true’ MAFs were beta-distributed along the genome, and the estimated minor allele count at each site was drawn from a binomial distribution with number of trials equal to the read depth at that site, and probability equal to the “true” MAF at that site. **D**: Comparison of MAFs from 27 replicate pairs resequenced from RNA. The plot represents all MAF frequency comparisons for the 27 samples where both replicates had >50,000 unique mapped reads, limited to genomic sites where the MAF > 0.02 in at least one of the 54 replicates, and excluding sites observed to be variant in more than 20 samples from our whole dataset at MAF > 0.03. The green lines are the threshold value of 0.03.

### Within-host variant sites are present in the majority of SARS-CoV-2 samples

Amongst the iSNV sites we identified, most were only observed in one or two samples (Fig. 2A). However, the majority of samples (305/462 with >50,000 unique reads) had at least one iSNV (Fig. 2B), consistent with previously reported levels (*13*). Two samples had a particularly high number (15 and 18) of iSNVs, each with high and correlated MAFs consistent with co-infection by two diverse variant strains (*14*). For one of these samples, laboratory contamination is unlikely since we could not identify any samples that could be the source, and independent epidemiological data is consistent with possible co-infection in this individual. We could not distinguish between co-infection and contamination in the other sample since both variant strains within it represent common genotypes in our study.

In general, the low level of genetic diversity of the virus makes identifying co-infection or contamination, and distinguishing between them, difficult. If sites where a large number of iSNVs are present are only observed to be variant within-host due to co-infection or contamination, then we estimate between ~1 to 2% of samples are potentially affected by co-infection or contamination (Table S2). As a precaution against contamination or batch effects, we sequenced known epidemiologically linked samples in different batches where possible (Fig. S3).

We hypothesised that a proportion of observed within-host variation could be due to co-infection with seasonal coronaviruses, which has been reported in 1-4% of SARS-CoV-2 infections (*15, 16*). Specifically, closely-matching reads from similar viruses could be mapped to SARS-CoV-2 and appear as mixed base calls. To understand the impact of co-infection, we re-captured and analysed a random subset of 180 samples spanning the full range of observed SARS-CoV-2 VLs (Ct 14 to 33, median 19.8), using the Castanet multi-pathogen enrichment panel (*9*), which contains probes for all known human coronaviruses with the exception of SARS-CoV-2. Among the 111 samples that yielded both SARS-CoV-2 and Castanet data, we identified one sample that was also positive for another betacoronavirus, human coronavirus OC43 (Fig. S4). Within the SARS-CoV-2 genome from this sample, which was complete and high-depth, we observed only a single iSNV at position 28580 and no evidence of mixed base calls at any other genomic position. This suggests that even where co-infection is present, it does not impact on the estimation of within-host diversity in our protocol, and the observed intrahost variation is indeed evidence of evolution of SARS-CoV-2.

**Fig. 2.**
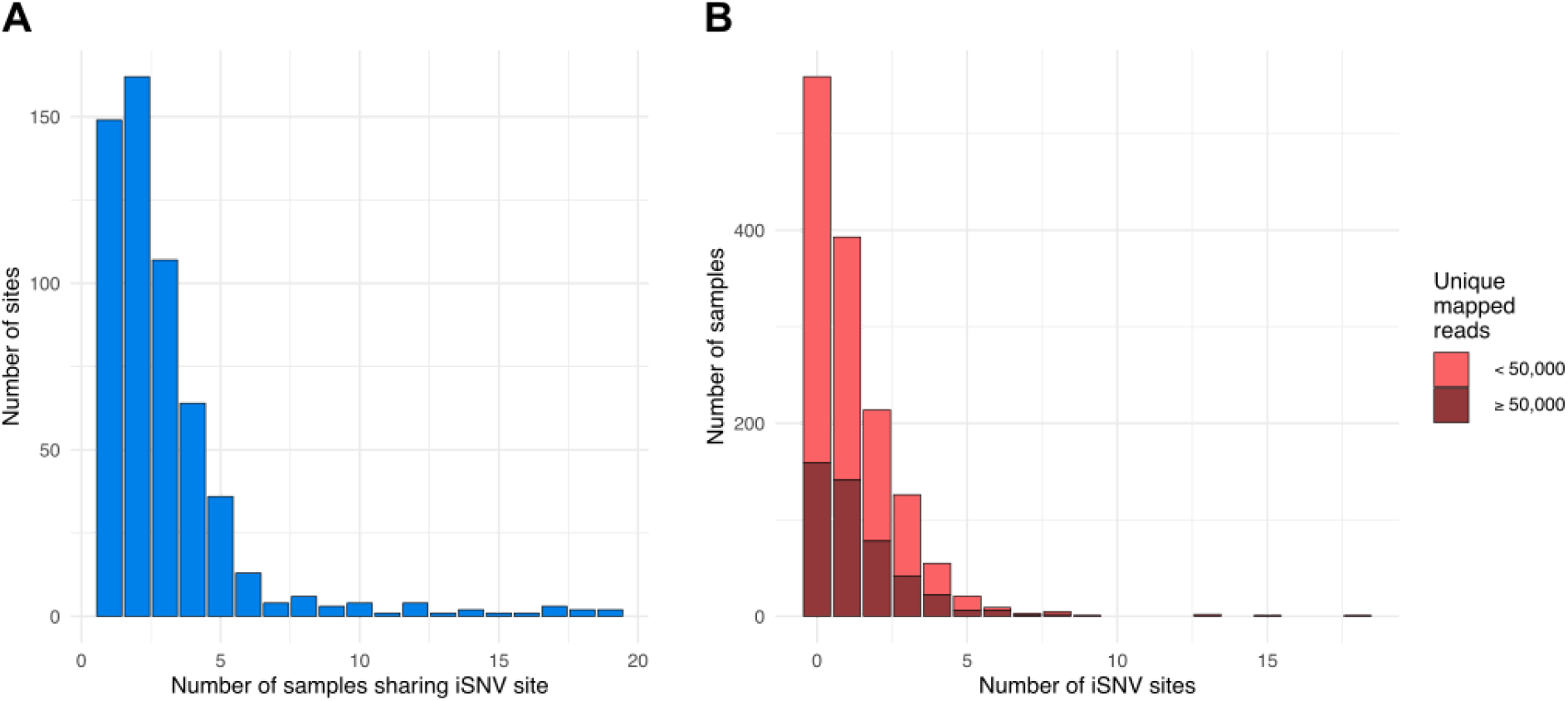
Intra-host variable (iSNV) sites are present in most samples and often shared. **A:** Distribution of the number of samples with an intra-host variant at a site. **B:** Distribution of the number of sites with iSNVs for all samples with more than 50,000 mapped reads (dark) and samples with fewer than 50,000 mapped reads (light). Only identified sites were included (see main text) with sites variable in 20+ individuals excluded.

### SARS-CoV-2 is evolutionary constrained at the within-host level

The distribution of iSNV sites varies across the genome (Table 1). Even excluding the UTR regions, which have a highly elevated density of iSNV sites, there is considerable variability across the genome, with open-reading frames (ORFs) 3a, 7a, and 8, and nucleocapsid (N) showing the highest densities. Most areas of the genome appear to be under strong purifying selection, with dn/ds values less than 1, including S. However, a few regions seem to be prone to directional selection within individuals, notably ORFs 3a, 7a, 7b, and 10. These patterns are broadly consistent with *dn/ds* values calculated for SNPs among consensus genomes (*17*), suggesting evolutionary forces at the within-host level are reflected at the between-host level, at least for within-host variant sites in high VL samples. The exception is ORF7a which appears to be under purifying selection at the between-host level, but positive selection at the within-host level.

**Table 1.** iSNVs and dn/ds by gene and over the whole genome.

| Gene | Start | End | Length | iSNVs |  |  | Mean iSNVs<br>per 100 sites | dn/ds |
| --- | --- | --- | --- | --- | --- | --- | --- | --- |
|  |  |  |  | Total | NS | S |  |  |
| 5'UTR | 1 | 265 | 265 | 40 | - | - | 0.0223 | - |
| ORF1a | 266 | 13483 | 13218 | 277 | 176 | 101 | 0.00311 | 0.43 |
| nsp1 | 266 | 805 | 540 | 17 | 9 | 8 | 0.00719 | 0.216 |
| nsp2 | 806 | 2719 | 1914 | 53 | 34 | 19 | 0.00395 | 0.422 |
| nsp3 | 2720 | 8554 | 5835 | 96 | 59 | 37 | 0.00216 | 0.433 |
| nsp4 | 8555 | 10054 | 1500 | 48 | 32 | 16 | 0.00484 | 0.452 |
| nsp5A | 10055 | 10972 | 918 | 13 | 12 | 1 | 0.00196 | 2.55 |
| nsp6 | 10973 | 11842 | 870 | 24 | 14 | 10 | 0.00513 | 0.35 |
| nsp7 | 11843 | 12091 | 249 | 4 | 2 | 2 | 0.00173 | 0.284 |
| nsp8 | 12092 | 12685 | 594 | 7 | 4 | 3 | 0.00157 | 0.361 |
| nsp9 | 12686 | 13024 | 339 | 10 | 6 | 4 | 0.00318 | 0.307 |
| nsp10 | 13025 | 13441 | 417 | 5 | 4 | 1 | 0.00276 | 1.99 |
| nsp12* | 13442 | 16236 | 2795 | 53 | 31 | 22 | 0.00314 | 0.333 |
| ORF1b | 13468 | 21555 | 8088 | 166 | 100 | 66 | 0.0031 | 0.374 |
| nsp13 | 16237 | 18039 | 1803 | 34 | 20 | 14 | 0.00235 | 0.442 |
| nsp14 | 18040 | 19620 | 1581 | 41 | 24 | 17 | 0.00419 | 0.267 |
| nsp15 | 19621 | 20658 | 1038 | 21 | 14 | 7 | 0.00215 | 0.649 |
| nsp16 | 20659 | 21552 | 894 | 17 | 11 | 6 | 0.00362 | 0.431 |
| S | 21563 | 25384 | 3822 | 84 | 55 | 29 | 0.00358 | 0.577 |
| ORF3a | 25393 | 26220 | 828 | 42 | 35 | 7 | 0.00938 | 1.43 |
| E | 26245 | 26472 | 228 | 6 | 4 | 2 | 0.0041 | 0.682 |
| M | 26523 | 27191 | 669 | 16 | 10 | 6 | 0.00344 | 0.443 |
| ORF6 | 27202 | 27387 | 186 | 6 | 5 | 1 | 0.00387 | 0.971 |
| ORF7a | 27394 | 27759 | 366 | 17 | 15 | 2 | 0.00806 | 1.47 |
| ORF7b | 27756 | 27887 | 132 | 5 | 5 | 0 | 0.00436 | ∞ |
| ORF8 | 27894 | 28259 | 366 | 18 | 10 | 8 | 0.00963 | 0.466 |
| N | 28274 | 29533 | 1260 | 70 | 49 | 21 | 0.00828 | 0.577 |
| ORF10 | 29558 | 29674 | 117 | 4 | 3 | 1 | 0.00676 | 1.31 |
| 3'UTR | 29675 | 29903 | 229 | 26 | - | - | 0.0232 | - |
| All coding positions** | 266 | 29674 | 29256 | 709 | 465 | 244 | 0.0038 | 0.492 |
| Full genome | 1 | 29903 | 29903 | 781 | - | - | 0.00411 | - |

All genome positions are relative to the Wuhan-Hu-1 reference sequence. iSNVs at the 18 “highly shared” sites and those identified from the synthetic controls are excluded, as are those in the poly-A tail (positions 29865-29903). The “mean iSNVs per 100 sites” column is the mean number in each gene over all 1390 samples. Note that due to gene overlap and non-coding intergenic regions, the total number of iSNVs (781) cannot be obtained as the sum of any column in this table, even if the rows for nonstructural proteins in ORF1ab are excluded. * nsp12 overlaps the boundary between ORF1a and ORF1b. ** Intergenic regions are excluded from this row, for which the start and end points do not represent a continuous range.

### Within-host variant sites are phylogenetically associated

Consensus viral sequences that cluster closely on a phylogenetic tree have been used successfully in SARS-CoV-2 to identify epidemiological links (*18*–*20*). Due to the recent emergence and low evolutionary rate of SARS-CoV-2, its global phylogeny has only limited genetic diversity, and hence limited resolution to identify clusters. We sought to gain a better understanding of SARS-CoV-2 evolution and determine whether iSNVs could be used to help resolve phylogenies and transmission clusters. For the 1390 samples in our study, we constructed a phylogeny using the robust procedure outlined by (*21*) (Fig. 3A). Using this tree, we determined whether iSNVs, and SNPs (indicating a difference in the most common variant among samples) at the same locus, are phylogenetically associated.

For the 153 iSNVs that are also consensus SNPs in at least one sample, termed iSNV-SNPs, we examined the proximity of tips with the iSNV to the position of consensus changes (between the two most common bases at the site of the iSNV) on the phylogeny (see Methods). A highly significant negative association (one-sided Mann-Whitney U-test, *p* <3×10^-16^; Fig. S5A) was found between the presence of an iSNV at a given site in a sample and the patristic distance to the nearest example of a consensus change at the same site. When we tested sites individually, six showed a significant association after Benjamini-Hochberg correction (*p*<0.05), reducing to five if only one sample from each individual was included.

In Fig. 3B we show the example of site 28580, with the red clade representing change from the global consensus G to A (a nonsynonymous change D103N in N), and nearby iSNVs occurring, both as minor As in the nodes ancestral to the change branch, and as minor Gs in the branch’s immediate descendants. Based on corresponding epidemiological data, this represents a likely healthcare-associated cluster with onward transmission to close contacts. In Fig. 3C we give the further example of site 20796, a synonymous substitution L6843 in ORF1a. Trees for the other significant sites after Benjamini-Hochberg correction appear in Fig. S6. In addition, we examined 16 epidemiologically identified household clusters, in 5 of which we observed an iSNV in one individual that was fixed in the other (for the household analyses we did not constrain on sites only present in high VL samples; Table 2).

For the 261 iSNVs that are present in at least two individuals but never reach consensus, we analysed the association with the phylogeny of each iSNV variant as a discrete trait, using two statistics: the association index (*22*) and the mean patristic distance between iSNV tips. After adjustment for multiple testing, no sites showed a *p-*value less than 0.05 for a phylogeny-iSNV association for either statistic. Similarly, if we simply compare the distance to the nearest iSNV tip amongst iSNV and non-iSNV tips across all 261 iSNV sites, there is also no evidence for an association (one-sided Mann-Whitney U-test p~=1, Fig. S5B). Nevertheless, some individual sites do show patterns suggestive of iSNV transmission, with diversity maintained after transmission (22 with *p*<0.05 before adjustment for multiple testing for at least one of the two statistics; those 9 with *p*<0.025 are shown in Fig. S6) suggesting we may lack the power to statistically detect some associations. Among the 16 known household clusters, we observed only one iSNV shared in two individuals within the same household. This iSNV was unique to these two individuals, demonstrating a likely example of transmitted viral diversity (Table 2).

**Fig. 3.**
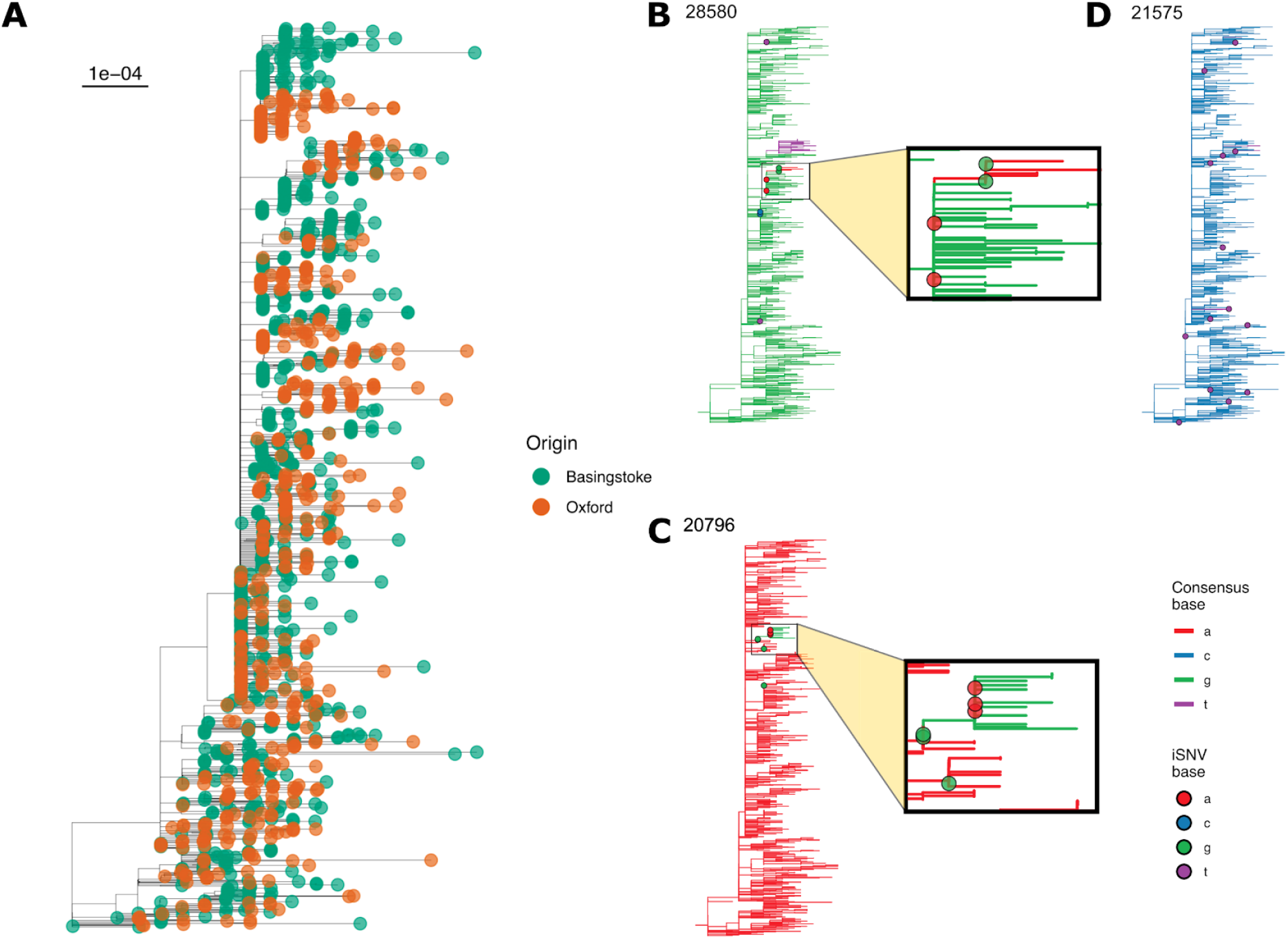
Consensus phylogeny of all isolates. In **A,** tips are coloured by sampling centre (Oxford or Basingstoke). The tree scale is in substitutions per site. Panels **B**-**D**: distribution of samples with iSNVs at three loci. The genomic coordinate (with respect to the Wuhan-Hu-1 reference sequence) appears in the top left. Tree branches are coloured by the consensus base at that position, and filled circles indicate samples iSNVs present at minimum 3% for samples with depth of at least 100 at that position, and are coloured by the most common minor variant present. For sites 28580 (**B**) and 20796 (**C**), an inset panel enlarges the a section of the phylogeny where a consensus change is in close proximity to iSNVs with the relevant pair of nucleotides involved.

### The transmission bottleneck size within households is small

Estimating bottleneck size is difficult for SARS-CoV-2, since it requires sufficient genetic diversity to differentiate distinct viruses that may be transmitted in known source-recipient pairs, and confidence that transmission is the cause of variants observed in both source and recipients (*23*–*25*). Using the exact beta-binomial method (*23*) we estimated bottleneck sizes between 1 and 8 among 14 household transmission pairs (Table 2). These observations are consistent with the small bottleneck sizes observed for influenza (*25*).

We speculate that situations where multiple phylogenetically linked cases share sub-consensus variants could be a consequence of superspreader events, or other high-exposure situations, where many individuals are exposed to high viral doses. An association between the route of exposure and the transmission bottleneck has been demonstrated experimentally for influenza (*26*). Here, we sequenced clinical samples, which likely include infections from some high-exposure events. For example, the clearest example of shared diversity is at site 28580, with three individuals attending the same hospital department on the same day, and estimated bottleneck sizes of 4 between the assumed source (determined by date of positive test) and each of the two recipients.

Taken together, our observations suggest the transmission bottleneck can be wide enough to permit co-transmission of multiple genotypes in some instances, but small enough that multiple variants do not persist after a small number of subsequent transmissions. In the cases where this transmission culminates in a consensus change on the phylogeny these patterns are readily observable, but in most cases patterns of co-transmission are drowned out by the high proportion of iSNVs that fail to transmit, or are transmitted but then lost.

**Table 2.** Household analysis of variants and transmission bottleneck size.

| Household<sup>1</sup> | iSNV-consensus<sup>2</sup> | iSNV-iSNV<sup>3</sup> | Bottleneck size<sup>4</sup> |
| --- | --- | --- | --- |
| 1 | 0 | 0 | 3 |
| 2 | 0 | 0 | 1 |
| 3 | 0 | 0 | 2 |
| 4 | 0 | 1 | 5 |
| 5 | 1 | 0 | 1,2 |
| 6 | 0 | 0 | 1 |
| 7 | 0 | 0 | 1 |
| 8 | 0 | 0 | - |
| 9 | 0 | 0 | 1 |
| 10 | 0 | 0 | 1 |
| 11 | 0 | 0 | 5,8 |
| 12 | 1 | 0 | 1,2 |
| 13 | 2 | 0 | 2 |
| 14 | 1 | 0 | 1 |
| 15 | 6 | 0 | - |
| 16 | 0 | 0 | 1,6 |

<sup>1</sup>All households consisted of two individuals with sequence data.
<sup>2</sup>The number of genome positions where a minor variant in one of the individuals (defined as >3% MAF and more than 100 reads) is the consensus variant in the other individual in the household (in all cases the consensus variant was >99.5%). All (non-masked) genomic sites were considered.
<sup>3</sup>The number of genome positions where a minor variant is >3% MAF in both individuals in the household.
<sup>4</sup>Bottleneck size was calculated using the exact beta-binomial method described in (23). All sites >3% MAF and more than 100 reads in the assumed source individual were used in the analysis. In the recipient all reads at these sites were considered, with an error threshold of 0.5% MAF. Where the first samples for each individual in the household were more than one week apart, we assumed the earlier sampled individual was the donor. Where an individual had more than one sample, and/or the first individuals were positively sampled within a week, we calculated all possible combinations of donor-recipient samples. The maximum and minimum maximum likelihood estimates are recorded if different. No estimate is recorded if there are no identified iSNVs >3% in the donor (household 8), or the two individuals in the household had more than two consensus differences (household 15).

### Some within-host variants show signatures of selection

Variants occurring repeatedly, but without phylogenetic association, could indicate sites under selection in distinct individuals (*27*). Of particular note are variants we observed at three sites in S: 21575 (V5F), 22899 (G446V) and 24198 (A879V), with G446V lying within the receptor binding domain (RBD). The minor variant F5 was observed in 14 samples, and represented SNPs in 8 samples, but did not have phylogenetic association in our iSNV-SNP analysis (*p* = 0.771 before multiple testing adjustment, Fig. 3D). This V5F mutation has been shown to increase infectivity *in vitro (28),* and has previously been identified as a potential site subject to selection (*29*). This variant has repeatedly been observed in global samples, including as minority variant, but appears to be increasing in frequency slowly if at all, suggesting it is only advantageous within a small subset of individuals, with the variant either ‘reverting’ in subsequent infections (as seen in HIV (*30*)), or failing to transmit at all. Similarly, we observed the minor variants V446 and V879 in 4 and 6 individuals respectively. Both variants have previously been shown to reduce sensitivity to convalescent sera *in vitro (28),* and V446 strongly reduces binding of one of the antibodies (REGN10987) in the REGN-Cov2 antibody cocktail (*31*), suggesting these may represent antibody escape mutations.

### Implications of intra-host variation on consensus phylogenies

The presence of minority variants could explain some phylogenetic inconsistencies observed in SARS-CoV-2. The global phylogeny is reconstructed from an alignment with relatively few SNPs, and therefore the particular base identified as consensus at iSNV sites can affect the overall phylogeny (*32*). Minority variants can result in changes to branch lengths, either by shortening branches due to lack of resolution at an informative site, or by extending branches if a minor variant - real or artifactual - is miscalled as consensus, as may occur in low VL samples. Miscalling a minor variant as consensus can also generate homoplasies (sites that are repeatedly mutated on the SARS-CoV-2 phylogeny), particularly where the minor variant was the result of contamination or co-infection and represents a lineage-defining SNP in another part of the phylogeny. The same effect would be expected for sites that are prone to host RNA editing or RNA degradation, resulting in the same minority variants arising in different parts of the phylogeny (*33*).

For the iSNV-SNP associated sites that we detected, representing the emergence and/or transmission of genuine variants, our phylogeny appears to be robust to the presence of iSNVs. We observed relatively few homoplasies on our tree (97 out of 1254 SNPs; Table S3), which suggests that at least in our dataset, the presence of minority variants did not strongly impact the phylogenetic signal. However, some of the longest terminal branch lengths in our phylogeny were indeed associated with low VL samples and high MAFs (Fig. S7), which suggests that in some cases, minor variants could be responsible for branch length extension in low VL samples. While the presence of high-MAF, consensus-impacting minor variants in such samples could be due to the effect of proportional sampling from a smaller viral population (Fig. 1A,C), genuine biological explanations are also plausible, including late infection being associated with both low VLs and higher diversity (*34*), or of an association of low VL with RNA degradation, host editing, or deleterious mutations which in turn reduce the likelihood of onwards transmission.

We emphasise however, that the presence of iSNVs at common SNP and/or homoplasic sites is not necessarily indicative of co-infection or contamination, or the generation of methodological variants. The generation and transmission of iSNVs is a prerequisite for the generation of SNPs on the phylogeny, and homoplasic sites may represent sites under diversifying selection (positively selected in some individuals but negatively selected in others), or sites prone to generation of within-host variants. Nonetheless, as is increasingly being recognised, care is needed when both calling iSNVs and SNPs (*32*). By sequencing synthetic RNA controls, resequencing samples, and only identifying sites if variable in at least one high VL sample, we retained only high confidence variants for our analyses.

### Concluding remarks

We uncovered a consistent and reproducible pattern of within-host SARS-CoV-2 diversity in a large dataset of over 1000 individuals, with iSNV sites showing strong phylogenetic clustering patterns if they are also associated with a change in the consensus variant at the same site. However, most samples harboured few variant sites, with a pattern of strong within-host evolutionary constraint in most regions of the genome, including Spike. This indicates that the within-host emergence of vaccine- and therapeutic-escape mutations is likely to be relatively rare. Moreover, the transmission bottleneck size was very small (between 1 and 8) in most instances where we had epidemiological data, suggesting that even if escape-mutations do arise they will be prone to loss at the point of transmission.

Although this bodes well for the longevity of vaccines and antibody-based treatments, we observed two mutations in Spike (G446V and A879V) that have previously been shown to escape antibody binding (*28, 31*), and a third that has been shown to increase viral infectivity (V5F, (*31*)), emphasising the need for continuing vigilance. We identified 30 nonysynonymous iSNVs in Spike that are present in multiple individuals (Table S2), and we suggest these and other commonly occurring iSNVs in other regions of the genome should be investigated and monitored, particularly as vaccines and therapeutics are rolled out more widely.

Throughout, we aimed to minimise sequencing artefacts and sample contamination where possible. The dense sampling and deep sequencing of SARS-CoV-2 has enabled us to witness ‘evolution-in-action’, with diversity generated in one individual leading to a change in consensus and fixation in subsequently infected individuals. The observation of shared diversity among phylogenetically and epidemiologically linked individuals suggests within-host variants could be used, at least in some instances, to help better resolve patterns of transmission in a background of low consensus diversity.

Our work demonstrates that an essential requirement for incorporating intrahost variants in any analysis is an understanding of the population prevalence of intrahost diversity, conditional on the methods used to produce the deep sequencing data. Moreover, our results emphasise the power of open data, large and rigorously controlled datasets, and the importance of integrating genomic, clinical, and epidemiological information, to gain in depth understanding of SARS-CoV-2 as the pandemic unfolds.

## Data and materials availability

All genomic data has been made publicly available as part of the COVID-19 Genomics UK (COG-UK) Consortium (*35*) via GISAID (*36*) and via the European Nucleotide Archive (ENA) study PRJEB37886.

## Supplementary Materials

Material and methods

Figures S1-S8

Tables S1-S3

OVSG Analysis group Membership

COG-UK full list of consortium names and affiliations

## Material and methods

Figures S1-S8

Tables S1-S3

OVSG Analysis group Membership

COG-UK full list of consortium names and affiliations

## Materials and methods

### RNA extraction

Residual RNA from COVID-19 RT-qPCR-based testing was obtained from Oxford University Hospitals (‘Oxford’), extracted on the QIASymphony platform with QIAsymphony DSP Virus/Pathogen Kit (QIAGEN), and from Basingstoke and North Hampshire Hospital (‘Basingstoke’), extracted with one of: Maxwell RSC Viral total nucleic acid kit (Promega); Reliaprep blood gDNA miniprep system (Promega); or Prepito NA body fluid kit (PerkinElmer). An internal extraction control was added to the lysis buffer prior to extraction to act as a control for extraction efficiency (genesig qRT-PCR kit, #Z-Path-2019-nCoV in Basingstoke, MS2 bacteriophage (*37*) in Oxford). The #Z-Path-2019-nCoV control is a linear, synthetic RNA target based on sequence from the rat *ptprn2* gene, which has no sequence similarity with SARS-CoV-2 (GENESIG PrimerDesign pers. comm, 6 April 2020). The MS2 RNA likewise has no SARS-CoV-2 similarity (*37*). Neither control RNA interfered with sequencing.

### Targeted metagenomic sequencing

Samples with suspected epidemiological linkage, where this information was available prior to sequencing, were processed in different batches. Sequencing libraries were constructed from remnant volume of nucleic acid after clinical testing, ranging from 5 to 45 μl (median 30μl) for each sample depending on the available amount of eluate. These volumes represented 1-15% of the original specimen (swab). Libraries were generated following the veSEQ protocol (*8*) with some modifications. Briefly, unique dual indexed (UDI) libraries for Illumina sequencing were constructed using the SMARTer Stranded Total RNA-Seq Kit v2—Pico Input Mammalian (Takara Bio USA, California, US) with no fragmentation of the RNA. An equal volume of library from each sample was pooled for capture. Size selection was performed on the captured pool to eliminate fragments shorter than 400nt, which otherwise may be preferentially amplified and sequenced. Target enrichment of SARS-CoV-2 libraries in the pool was obtained through a custom xGen Lockdown Probes panel (IDT, Coralville, USA), using the SeqCap EZ Accessory Kits v2 and SeqCap Hybridization and Wash Kit (Roche, Madison, US) for hybridization of the probes and removal of unbound DNA. Following 12 cycles of PCR for post-capture amplification, the final product was purified using Agencourt AMPure XP (Beckman Coulter, California, US). Sequencing was performed on the Illumina MiSeq (batches 1-2) or NovaSeq 6000 (batches 3-27) platform (Illumina, California, US) at the Oxford Genomics Centre (OGC), generating 150bp or 250bp paired-end reads.

### Quantification controls

A dilution series of *in vitro* transcribed SARS-CoV-2 RNA (Twist Synthetic SARS-CoV-2 RNA Control 1 (MT007544.1),Twist Bioscience) was included in every capture pool of 90 samples starting from batch 3, and sequenced alongside the clinical samples. Control RNA was serially diluted into Universal Human Reference RNA (UHRR) to a final concentration of SARS-CoV-2 RNA of 500,000, 50,000, 5,000, 500, 100 and 0 copies/reaction. From this we produced a standard curve demonstrating linear association between viral load (VL) and read depth (Fig. S1). For an experiment comparing iSNV presence with and without probe capture, we additionally sequenced two replicates of the Twist RNA control without capture, diluted into UHRR to give an expected concentration of 50,000 copies per reaction.

As an additional validation step, we compared intrahost single-nucleotide variants (iSNVs) in re-sequenced controls with data for the stock RNA sequenced and provided by the manufacturer (Twist Bioscience). Six well-defined iSNVs, which were present in the manufacturer’s data and presumably arose during in vitro transcription, were also recovered by our protocol (Fig. S8). In addition, we identified 112 sites that appeared vulnerable to low-frequency intrahost variation in vitro (Table S3), possibly as a result of structural variation along the genome or interaction with the sequencing protocol. We blacklisted vulnerable sites from further analysis.

### In-run controls

In addition to the synthetic RNA standards described above, each batch included a non-SARS-CoV-2 in-run control consisting of purified *in vitro* transcribed HIV RNA from clone p92BR025.8, obtained from the National Institute for Biological Standards and Control (NIBSC) (*38*). For batches 1 and 2, which were sequenced prior to synthetic RNA becoming available, we included negative buffer controls. As additional negative controls, we sequenced 6 matched clinical samples from non-COVID-19 patients, distributed across different sequencing runs; none contained any SARS-CoV-2 reads.

### Minimising risk of index misassignment

All samples had unique dual indexing (UDI) to prevent cross-detection of reads in the same pool. We used the in-run HIV RNA controls to estimate index misassignment, as this provided a sequence-distinct source of RNA: <3 SARS-CoV-2 reads were detected in any HIV control (median 0) and <10 HIV reads were detected in any SARS-CoV-2 control (median 0), suggesting that index misassignment, if present, occurred at extremely low levels.

### Bioinformatics processing

De-multiplexed sequence read pairs were classified by Kraken v2 (*39*) using a custom database containing the human genome (GRCh38 build) and the full RefSeq set of bacterial and viral genomes (pulled May 2020). Sequences identified as either human or bacterial were removed using filter_keep_reads.py from the Castanet (*9*) workflow (https://github.com/tgolubch/castanet). Remaining reads, comprised of viral and unclassified reads, were trimmed in two stages: first to remove the random hexamer primers from the forward read and SMARTer TSO from the reverse read, and then to remove Illumina adapter sequences using Trimmomatic v0.36(*40*), with the ILLUMINACLIP options set to “2:10:7:1:true MINLEN:80”. Trimmed reads were mapped to the SARS-CoV-2 RefSeq genome of isolate Wuhan-Hu-1 (NC_045512.2), using shiver (*41*) v1.5.7, with either smalt(*42*) or bowtie2 (*43*) as the mapper. Both mappers generated comparable results; smalt was used for the final analysis. Only properly paired reads with insert size under 2000 and with at least 70% sequence identity to the reference were retained. For analysis of consensus genomes, consensus calls required a minimum of 2 uniquely mapped (deduplicated) reads per position, equivalent to >15 raw reads per position. Analysis of within-host diversity was restricted only to positions with minimum raw depth of 100, except when examining diversity within presumed recipients of transmissions in the bottleneck analysis. Minor allele frequencies were computed at every position using shiver (*41*) (tools/AnalysePileup.py), with the default settings of no BAQ and maximum pileup depth of 1000000. Lineages were assigned by the Pangolin web server (https://pangolin.cog-uk.io) using the determined consensus genome for each sequenced sample.

### Alignment

Oxford and Basingstoke samples were selected if the consensus sequence (inferred from unique mapped reads) consisted of no more than 25% N characters. As an alignment to the reference sequence was already performed in *shiver,* no further alignment was necessary. To place these data into the global phylogenetic context and help resolve ancestry, a collection of non-UK consensus sequences from the GISAID database (*44*) were included in the set of sequences to be aligned. All GISAID (*36*) sequences were downloaded from the database on the 26th April 2020 and filtered to remove sequences that were less than 29800 base pairs in length, were more than 1% Ns, or were from the United Kingdom. The remaining sequences were clustered using CD-HIT-EST (*45*) using a similarity threshold of 0.995, and then one sequence per cluster picked. The resulting set, along with the reference genome Wuhan-Hu-1 (RefSeq ID NC_045512), were aligned using MAFFT (*46*), with some manual improvement of the algorithmic alignment and removal of problematic sequences performed as a post-processing step. Indels with respect to Wuhan-Hu-1 in both the Oxford/Basingstoke and GISAID alignments were deleted, resulting in two alignments of 29903 nucleotides that could be readily combined.

### Simulation of expected number of iSNVs for a given VL sample

To demonstrate the effect of read depth on estimated iSNV counts, we first assumed that within-host MAFs at each site *s* (here regarded as simply a proportion of reads that do not share the majority nucleotide at *s*) for each isolate *i* were drawn from a Beta(1, 331.41) distribution. Under this distribution, whose mode is 0, the expected number of sites across the whole genome with a MAF greater than 0.03 is 1.22, which is the mean number of iSNVs per sample in the real data. Let *p_si_* represent this “true” simulated MAF, and *d_si_* the empirical read depth, of sample *i* at site *s*. Then the estimated number of MAFs for *i* was calculated by summing draws from a Binomial(*d_s/_*, *p_si_*) distribution for each site *s*.

### Phylogenetics

Phylogenetic reconstruction was performed on the alignment consisting of the 1390 consensus sequences, along with the GISAID set and the Wuhan-Hu-1 reference sequence. We followed the recommendations of (*21*) whereby 100 separate maximum likelihood phylogenies were generated using RAxML-NG (*47*) and the GTR+G substitution model, such that each reconstruction used a different random starting parsimony tree. The final phylogeny was then obtained from this set using majority rule. This final tree was rooted with respect to the reference sequence, and then that and all GISAID isolates were pruned.

To identify homoplasic sites, we selected sites that changed state more than once along the tree, after inferring the states at internal nodes using ancestral state reconstruction as implemented in ClonalFrameML (*48*) and rooting the tree using the reference genome NC_045512.

### Calculation of *dn/ds*

The total number of synonymous and nonsynonymous substitutions in the SARS-CoV-2 genome was estimated using the first method of (*49*) applied to the coding regions of the Wuhan-Hu-1 reference sequence. Overlapping reading frames were accounted for such that a substitution was considered nonsynonymous overall if it was nonsynonymous in either frame. The d*n*/d*s* ratio for iSNVs over a genomic region *G* was then calculated as:

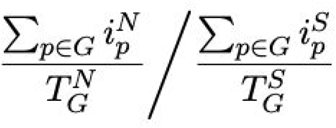

where 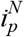 is the fraction of iSNVs at *p* that are nonsynonymous, or 0 if there are no iSNVs at *p*, 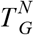 the total number of potential nonsynonymous substitutions in *G*, and the denominator replaces *N* with *S* to represent synonymous substitutions.

### Phylogenetic association of iSNVs and SNPs

Where an iSNV corresponded to a consensus SNP (by the base pair involved, not simply the site), we performed ancestral state reconstruction on the consensus trees using ClonalFrameML (*48*) to identify all branches upon which that substitution was involved. Tips derived from the same clinical sample were then pruned until only one (the one with the highest overall depth) remained. We then, for each tip in the tree, calculated the patristic distance from that tip to the midpoint of the closest one of these branches, and used a one-tailed Mann-Whitney U-test to test for association between the iSNV existing in a sample and this distance. Multiple testing was controlled for using the Benjamini-Hochberg adjustment. As a sensitivity analysis, this was repeated such that all but one tip per infected individual, rather than per clinical sample, were pruned. These analyses were done both on an individual site level and across all sites of interest.

### Phylogenetic association of iSNVs at consensus invariant positions

For the remaining iSNVs, we calculated the extent of association with the consensus phylogeny by treating the presence of an iSNV as a discrete character and calculating the association index, and the mean patristic distance between iSNV tips. Once again the consensus tree was pruned such that tips corresponding to samples with read depth <100 at the position and all but one tip coming from the same individual were removed. A null distribution was generated by permuting the tip labels of this tree 10,000 times, and a one-sided permutation test *p*-value calculated. Multiple testing was adjusted for as above. In addition, for each tip in the phylogeny at each site of interest, we calculated the minimum patristic distance to a different tip corresponding to an iSNV, and used the Mann-Whitney U-test again to compare the distribution of these distances between iSNV and non-iSNV tips.

### Figure S1-S8

**Supplementary Figure 1.**
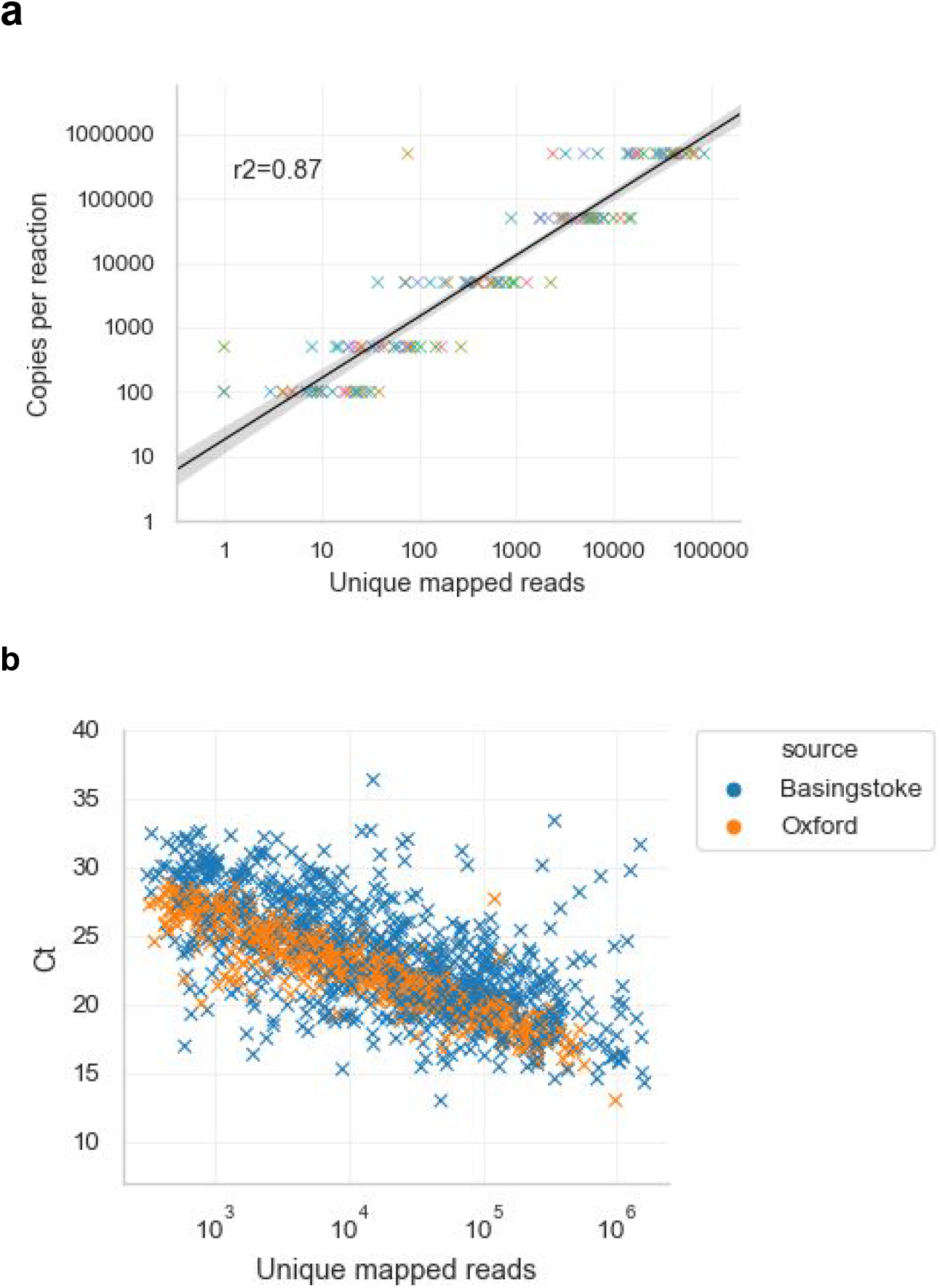
(a) Correlation between number of SARS-CoV-2 unique reads and RNA copies/ml for within-batch standard curves for dilution series of positive control RNA. Colour indicates batch. Synthetic SARS-CoV-2 RNA (generated by in vitro transcription by Twist Bioscience) was serially diluted into Universal Human Reference RNA (UHRR) to a final concentration of SARS-CoV-2 RNA of 500,000, 50,000, 5,000, 500, 100 and 0 copies/reaction. Controls were processed and sequenced alongside each batch of samples (batches 3-27). Batches 1 and 2 were processed prior to controls being available and did not have a standard curve. (b) Correlation between nearest available cycle threshold (Ct) value for sequenced clinical samples, as reported by the collecting laboratory, and the number of unique mapped reads. Due to variation in qPCR methodology, Ct values varied substantially between laboratories and over time. Higher RNA volumes were made available for sequencing for Basingstoke samples, contributing to the observation of higher read unique numbers (viral load) for the same Ct values, compared with Oxford samples.

**Supplementary Figure 2.**
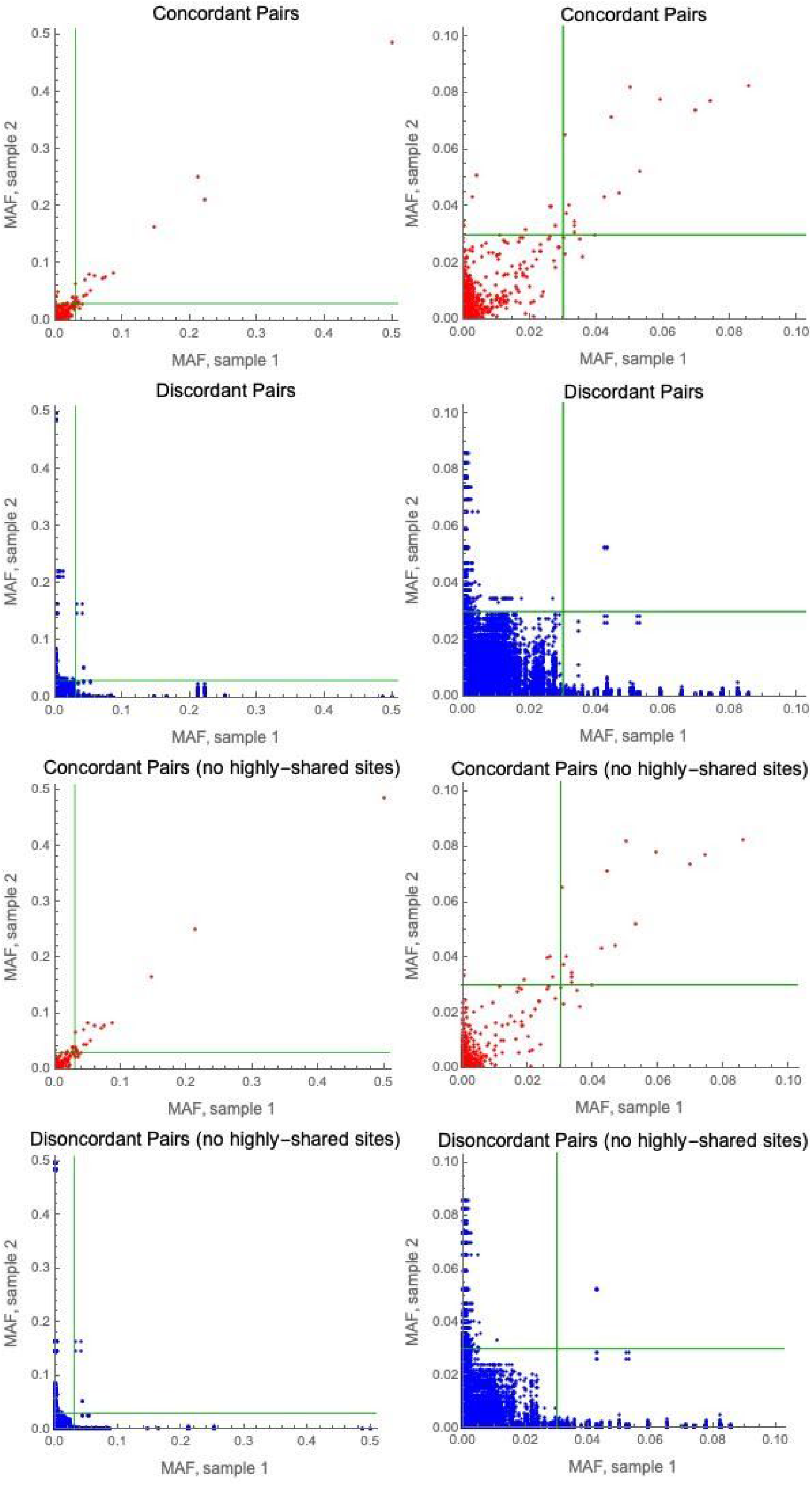
Comparison of minor allele frequencies among replicate samples. Data is only included for the 27 replicate pairs where both replicates had more than 50,000 unique mapped reads, and for all sites where MAF >=2% and depth >= 100 in at least one of the 54 replicates. For MAFs > 3%, and excluding highly-shared sites, MAFs are highly reproducible. Concordant pairs: The points represent the MAFs in each of the replicate pairs, for all 27 replicate pairs for all identified sites. If MAFs are reproducible, we expect a positive correlation. Discordant pairs: The points represent the MAFs for all pairwise permutations of replicates for all identified sites, excluding concordant replicates. Unless variants are present in multiple pairs of samples, the expectation is for points to be positioned along the axes. Top two rows include all sites, whereas the bottom two rows exclude highly-shared sites (those observed at MAF >=3% in 20 or more samples across the entire dataset). The blue points in the upper-right quadrants represent site 28580, which is present in phylogenetically linked individuals, with two of these included in the 27 replicate pairs. The green line shows MAF 3%.

**Supplementary Figure 3.**
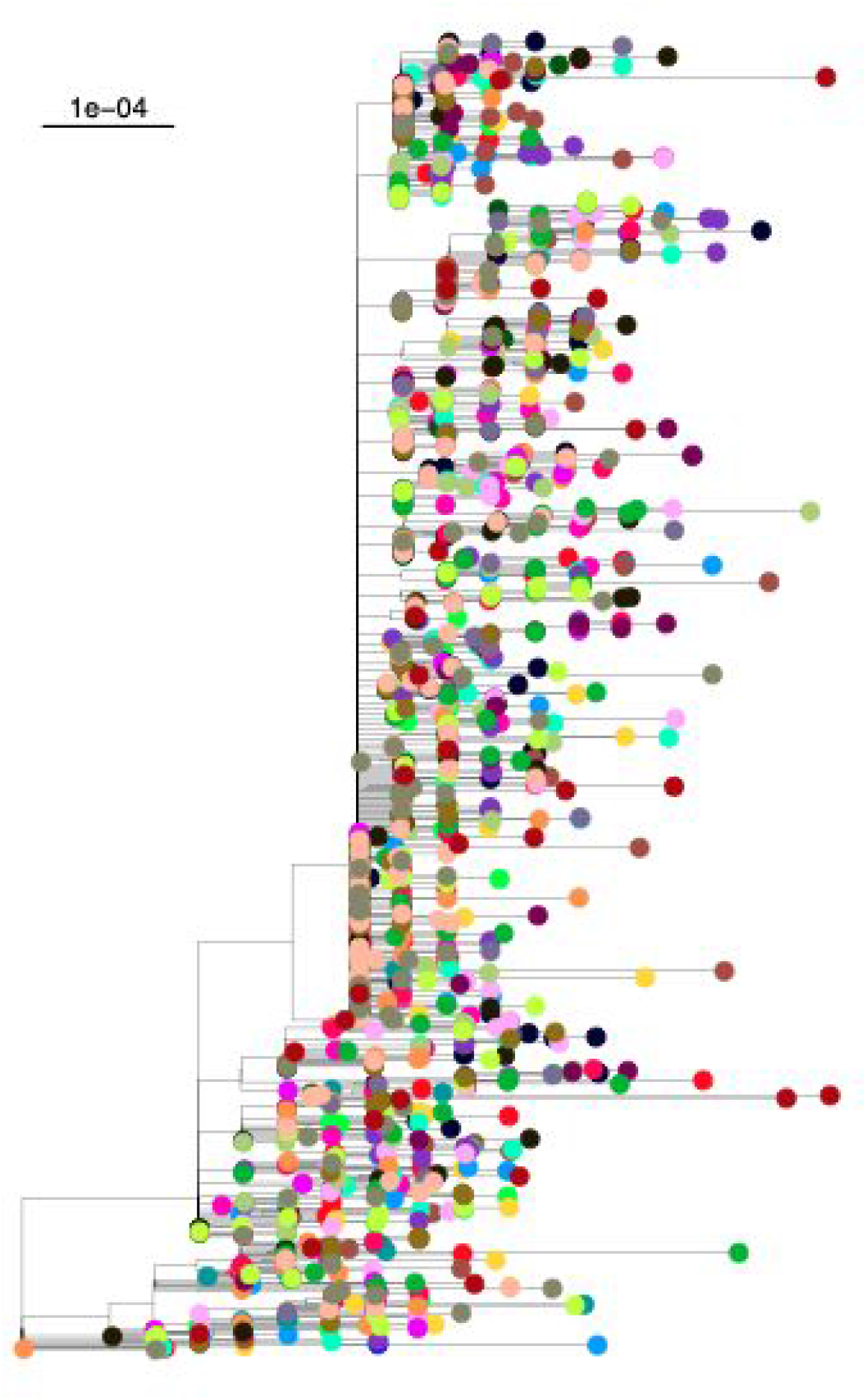
Consensus phylogeny of all 1390 Oxford and Basingstoke samples. Tips are coloured by sequencing batch.

**Supplementary Figure 4.**
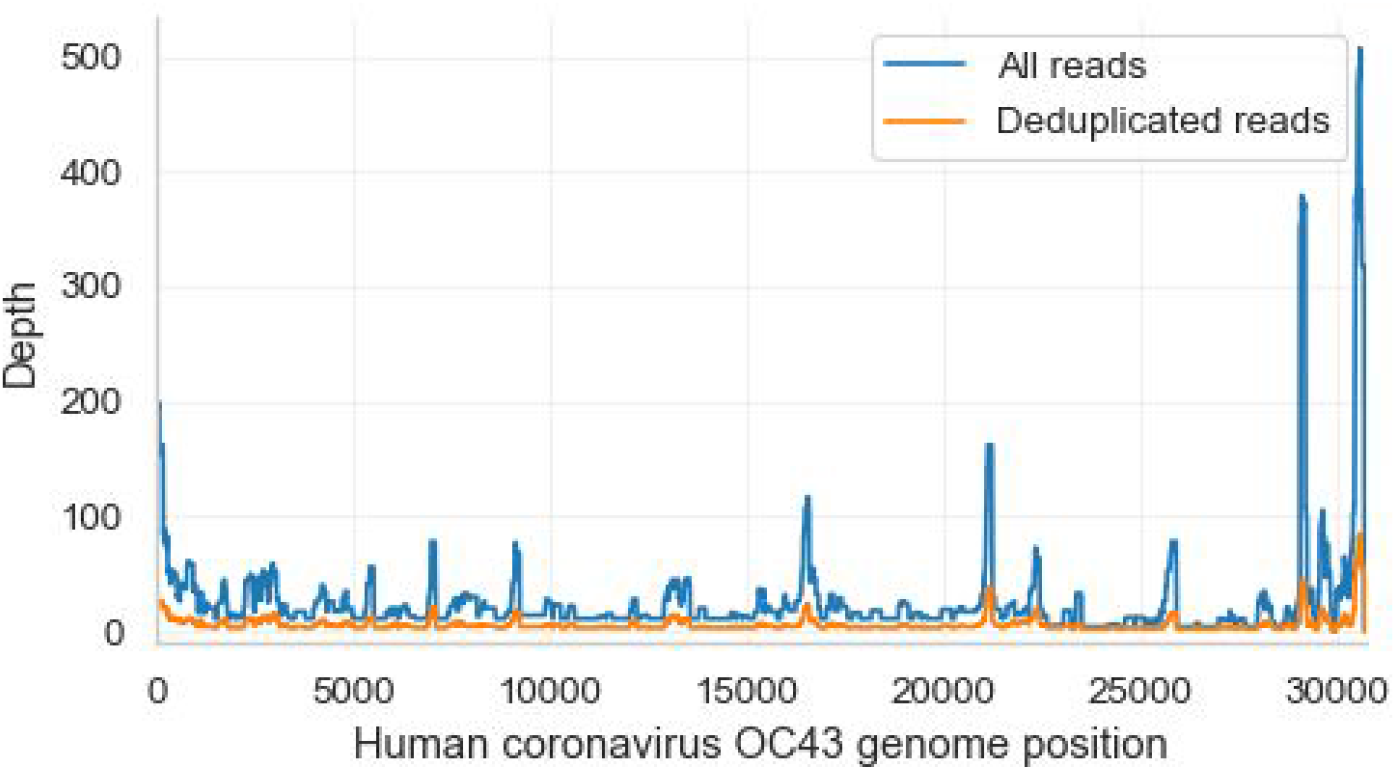
Genome coverage for co-infecting human coronavirus OC43 in a SARS-CoV-2-positive sample, OXON-AEC3D. A single co-infection with a non-SARS-CoV-2 circulating coronavirus was detected among a subset of 111 samples analysed with both SARS-CoV-2-specific probes and the Castanet metagenomic respiratory probe panel. Shown in blue are positions of the 2953 proper read pairs mapping to the Castanet reference for OC43, with unique (deduplicated) read depth in orange.

**Supplementary Figure 5.**
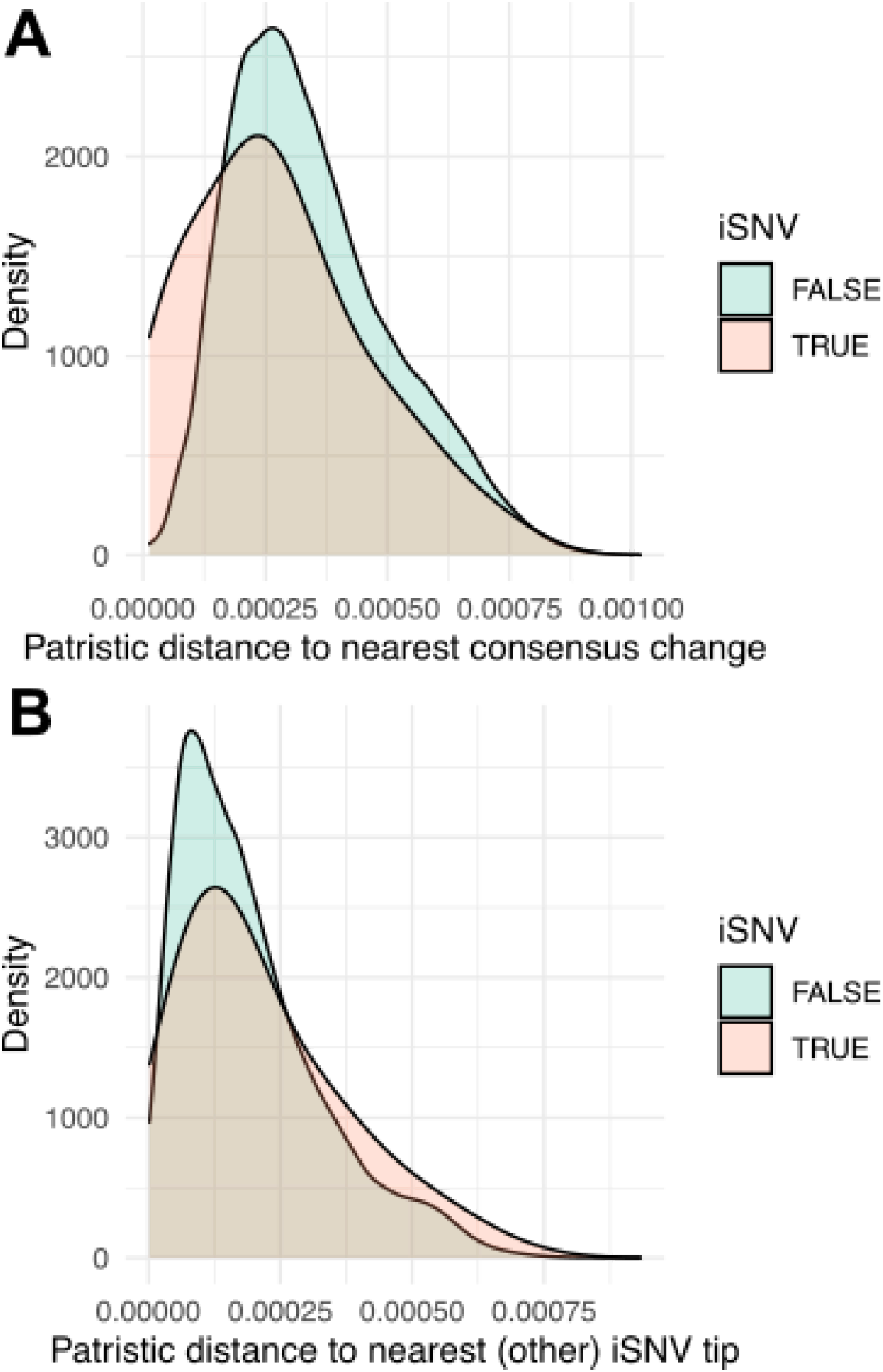
**A** Across all iSNVs that reach consensus, kernel density plot of the patristic distances from iSNV tips (orange) and other tips (green) to the nearest consensus branch change of the nucleotides involved. **B** Across all iSNVs that do not reach consensus and occur at least twice, kernel density plot of the patristic distances from iSNV tips and other tips to the nearest iSNV tip (other than the tip itself).

**Supplementary Figure 6.**
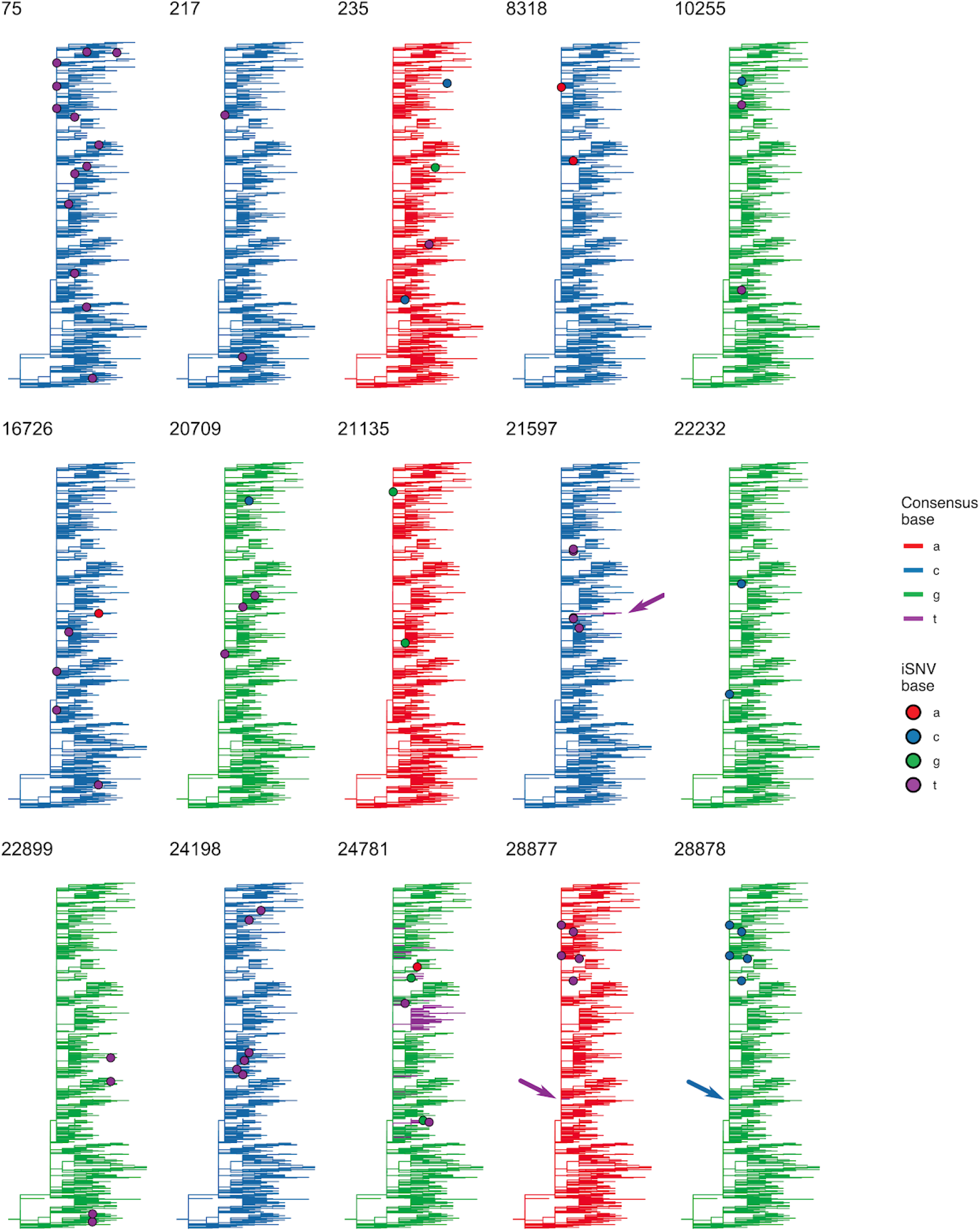
The consensus phylogeny coloured by SNP and iSNV for sites mentioned in the main text. Where consensus changes are hard to see they are indicated with an appropriately coloured arrow. Sites 21597, 24751, 28877 and 28878 are the remaining positions where a statistically significant association of iSNV tips with branches with a consensus base change was identified (along with 20796 and 28580). For some of these, coloured arrows indicate the presence of branches with consensus SNPs where this is difficult to see. Sites 22899 and 24198 are Spike variants shown to exhibit reduced sensitivity to convalescent sera (Li et al). The remaining 9 subfigures are for iSNVs which never reach consensus but show a *p*<0.025 for phylogenetic association of iSNV tips using the association index or the mean patristic distance between iSNV tips. While we lack the power to identify these once the Benjamini-Hochberg adjustment is applied, the patterns remain suggestive of transmission of iSNVs by eye.

**Supplementary Figure 7.**
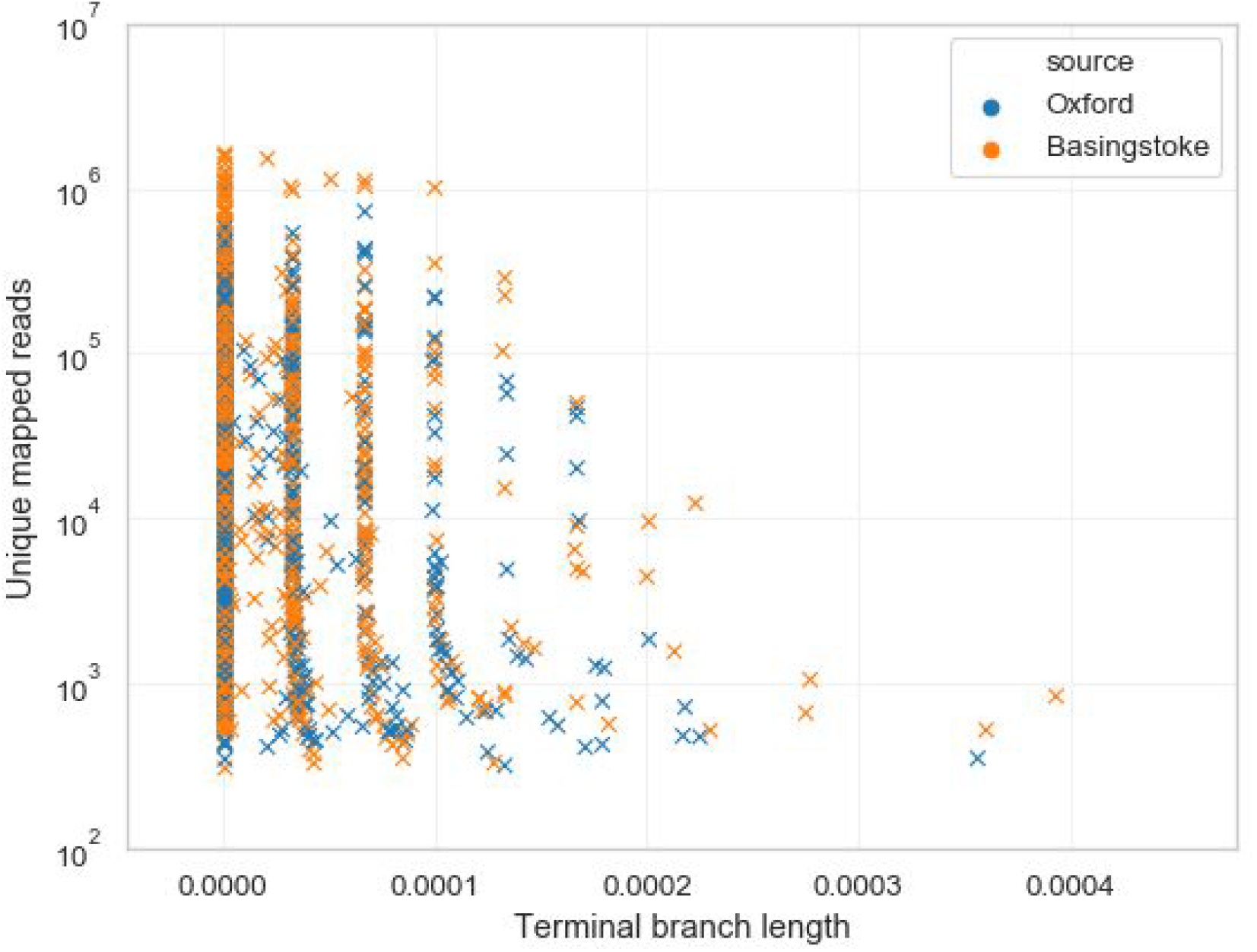
Relationship between terminal branch length and number of unique mapped reads (viral load indicator). Terminal branch lengths on the phylogeny of 1390 samples were plotted against the viral load as estimated from the number of uniquely mapped reads for each sample.

**Supplementary Figure 8.**
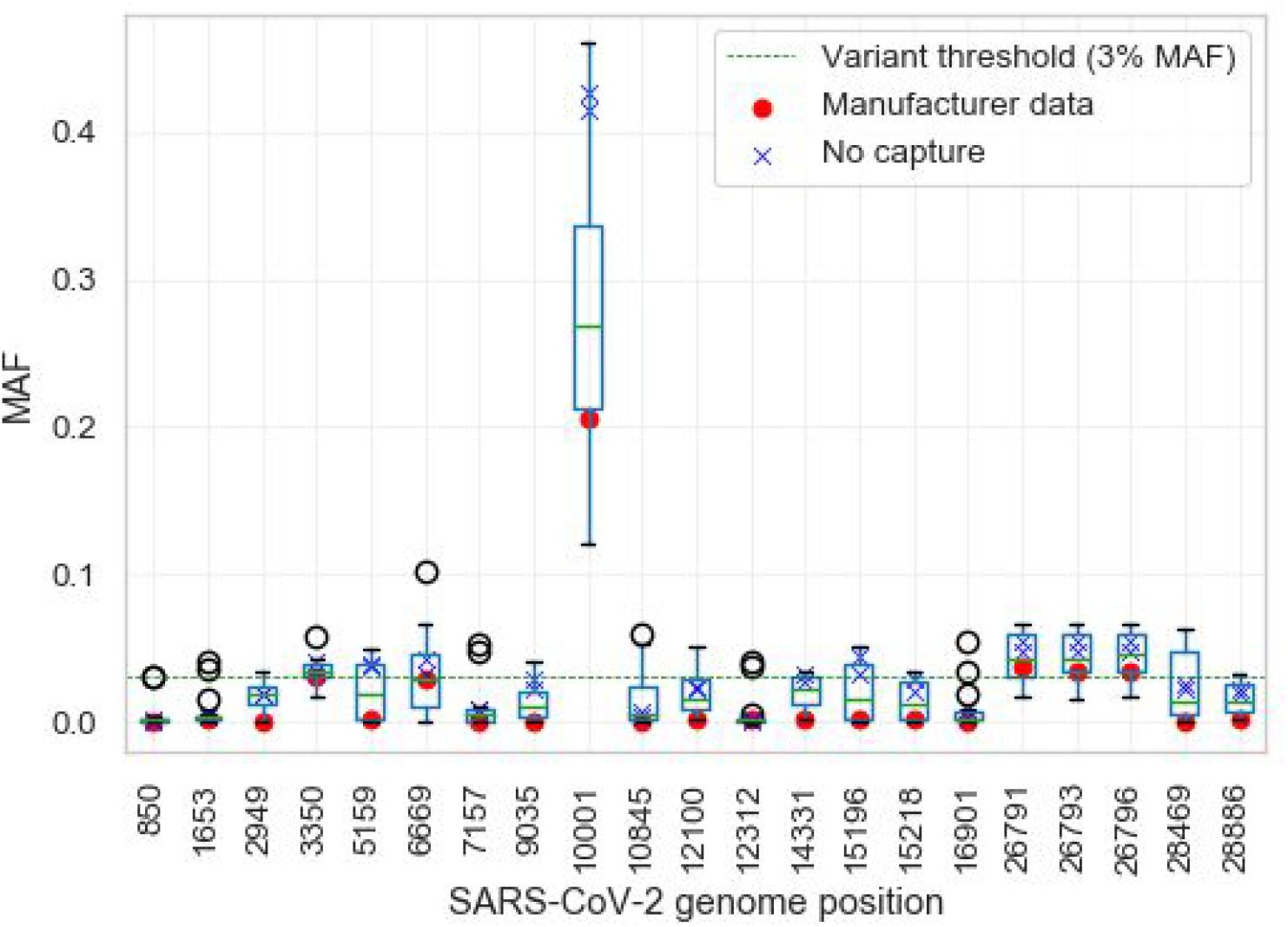
Within-sample diversity assessed in control RNA (Twist Bioscience). Within-sample diversity was assessed in RNA controls sequenced with each sequencing batch (0.5 mln copies per reaction). At all sites where at least 2 replicates had a minor variant with minimum 3% MAF (boxplot), diversity was compared against a set of NGS reads obtained from Twist Bioscience for the ancestral stock of the *in vitro* transcribed RNA used in this study (red circles). Six variants were consistently recovered from both the manufacturer data and the in-batch controls, at positions 3350, 6669, 10001, 26791, 26793, 26796. To check whether the remaining within-host variants arose during the SMARTer library prep or during probe capture, we additionally resequenced two replicates of the Twist RNA without capture (blue crosses), by diluting neat RNA 50:50 v/v in Universal Human Reference RNA (UHRR) and taking a proportion for sequencing, to yield approximately 50,000 copies of the Twist control RNA per sample. We generated SMARTer libraries from these replicates, and sequenced these alongside other samples in separate batches. The two capture-free replicates had the same range of intra-sample variants as were observed in our routinely sequenced controls, implying that any differences from the manufacturer data cannot be explained by probe capture and must be the result of the SMARTer library protocol and/or stochastic variation between our laboratory aliquot and the ancestral RNA stock sequenced by Twist.

### Tables S1-S3

**Table S1.**
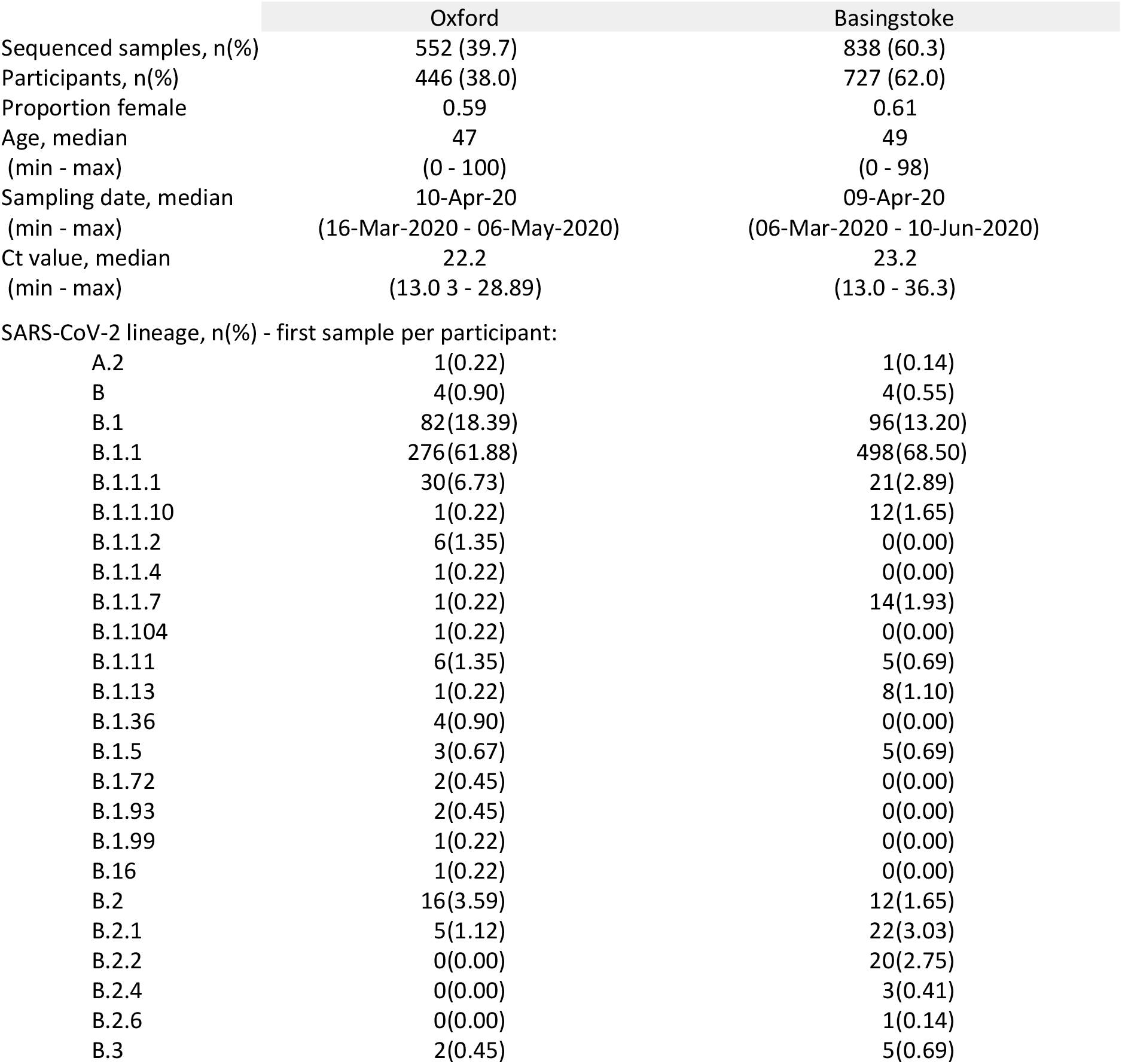
Baseline characteristics of SARS-CoV-2 samples in our dataset collected by participating hospitals in Oxford and Basingstoke, UK, between 8 March and 10 June 2020. Lineages are given for the first sample per participant, excluding anonymous samples.

**Table S2. Identified within-host variable sites.** Sites with at least one minor allele at frequency >= 3% at depth of at least 100 reads, in a sample depth >=50,000 unique mapped reads. Throughout, “samples” refers to all sequencing runs, and therefore includes replicates in the totals. n_notPopConsensus refers to the number of samples in which the minor variant is not the population-level consensus (most common consensus allele); n_SNPs gives the number of SNPs on the tree; homoplasy is “TRUE” if a homoplasy exists on the tree; maf_median is the median minor allele frequency (MAF) for all samples with >2% MAF; maf_IQR is the inter-quartile range of minor allele frequencies for all samples with >2% MAF. https://github.com/katrinalythgoe/COVIDdiversity

**Table S3. List of sites masked due to vulnerability to low frequency variation.** https://github.com/katrinalythgoe/COVIDdiversity

### OVSG Analysis Group membership

John A Todd, Tanya Golubchik, David Bonsall, Christophe Fraser, Derrick Crook, Tim Peto, Monique Andersson, Katie Jeffery, David Eyre, Timothy Walker, Robert Shaw, Peter Simmonds, Katrina Lythgoe, Luca Ferretti, Matthew Hall, Mariateresa de Cesare, Paolo Piazza, Richard Cornall.

**COG-UK Full list of consortium names and affiliations**

**Funding acquisition, leadership, supervision, metadata curation, project administration, samples, logistics, Sequencing, analysis, and Software and analysis tools:**

Thomas R Connor ^33, 34^, and Nicholas J Loman ^15^.

**Leadership, supervision, sequencing, analysis, funding acquisition, metadata curation, project administration, samples, logistics, and visualisation:**

Samuel C Robson ^68^.

**Leadership, supervision, project administration, visualisation, samples, logistics, metadata curation and software and analysis tools:**

Tanya Golubchik ^27^.

**Leadership, supervision, metadata curation, project administration, samples, logistics sequencing and analysis:**

M. Estee Torok ^8, 10^.

**Project administration, metadata curation, samples, logistics, sequencing, analysis, and software and analysis tools:**

William L Hamilton ^8, 10^.

**Leadership, supervision, samples logistics, project administration, funding acquisition sequencing and analysis:**

David Bonsall ^27^.

**Leadership and supervision, sequencing, analysis, funding acquisition, visualisation and software and analysis tools:**

Ali R Awan ^74^.

**Leadership and supervision, funding acquisition, sequencing, analysis, metadata curation, samples and logistics:**

Sally Corden^33^.

**Leadership supervision, sequencing analysis, samples, logistics, and metadata curation:**

Ian Goodfellow ^11^.

**Leadership, supervision, sequencing, analysis, samples, logistics, and Project administration:**

Darren L Smith ^60, 61^.

**Project administration, metadata curation, samples, logistics, sequencing and analysis:**

Martin D Curran ^14^, and Surendra Parmar ^14^.

**Samples, logistics, metadata curation, project administration sequencing and analysis:**

James G Shepherd ^21^.

**Sequencing, analysis, project administration, metadata curation and software and analysis tools:**

Matthew D Parker ^38^ and Dinesh Aggarwal ^1, 2, 3^.

**Leadership, supervision, funding acquisition, samples, logistics, and metadata curation:**

Catherine Moore ^33^.

**Leadership, supervision, metadata curation, samples, logistics, sequencing and analysis:**

Derek J Fairley^6, 88^, Matthew W Loose ^54^, and Joanne Watkins ^33^.

**Metadata curation, sequencing, analysis, leadership, supervision and software and analysis tools:**

Matthew Bull ^33^, and Sam Nicholls ^15^.

**Leadership, supervision, visualisation, sequencing, analysis and software and analysis tools:**

David M Aanensen ^1, 30^.

**Sequencing, analysis, samples, logistics, metadata curation, and visualisation:**

Sharon Glaysher ^70^.

**Metadata curation, sequencing, analysis, visualisation, software and analysis tools:**

Matthew Bashton ^60^, and Nicole Pacchiarini ^33^.

**Sequencing, analysis, visualisation, metadata curation, and software and analysis tools**: Anthony P Underwood ^1, 30^.

**Funding acquisition, leadership, supervision and project administration:**

Thushan I de Silva ^38^, and Dennis Wang ^38^.

**Project administration, samples, logistics, leadership and supervision**:

Monique Andersson^28^, Anoop J Chauhan ^70^, Mariateresa de Cesare ^26^, Catherine Ludden ^1,3^, and Tabitha W Mahungu ^91^.

**Sequencing, analysis, project administration and metadata curation:**

Rebecca Dewar ^20^, and Martin P McHugh ^20^.

**Samples, logistics, metadata curation and project administration:**

Natasha G Jesudason ^21^, Kathy K Li MBBCh ^21^, Rajiv N Shah ^21^, and Yusri Taha ^66^.

**Leadership, supervision, funding acquisition and metadata curation:**

Kate E Templeton ^20^.

**Leadership, supervision, funding acquisition, sequencing and analysis:**

Simon Cottrell ^33^, Justin O’Grady ^51^, Andrew Rambaut ^19^, and Colin P Smith^93^.

**Leadership, supervision, metadata curation, sequencing and analysis:**

Matthew T.G. Holden ^87^, and Emma C Thomson ^21^.

**Leadership, supervision, samples, logistics and metadata curation**:

Samuel Moses ^81, 82^.

**Sequencing, analysis, leadership, supervision, samples and logistics:**

Meera Chand ^7^, Chrystala Constantinidou ^71^, Alistair C Darby ^46^, Julian A Hiscox ^46^, Steve Paterson ^46^, and Meera Unnikrishnan ^71^.

**Sequencing, analysis, leadership and supervision and software and analysis tools:**

Andrew J Page ^51^, and Erik M Volz ^96^.

**Samples, logistics, sequencing, analysis and metadata curation:**

Charlotte J Houldcroft ^8^, Aminu S Jahun ^11^, James P McKenna ^88^, Luke W Meredith ^11^, Andrew Nelson ^61^, Sarojini Pandey ^72^, and Gregory R Young ^60^.

**Sequencing, analysis, metadata curation, and software and analysis tools:**

Anna Price ^34^, Sara Rey ^33^, Sunando Roy ^41^, Ben Temperton^49^, and Matthew Wyles ^38^.

**Sequencing, analysis, metadata curation and visualisation:**

Stefan Rooke^19^, and Sharif Shaaban ^87^.

**Visualisation, sequencing, analysis and software and analysis tools:**

Helen Adams ^35^, Yann Bourgeois ^69^, Katie F Loveson ^68^, Áine O’Toole ^19^, and Richard Stark ^71^.

**Project administration, leadership and supervision:**

Ewan M Harrison ^1 3^, David Heyburn ^33^, and Sharon J Peacock ^2, 3^

**Project administration and funding acquisition:**

David Buck ^26^, and Michaela John^36^

**Sequencing, analysis and project administration:**

Dorota Jamrozy ^1^, and Joshua Quick ^15^

**Samples, logistics, and project administration:**

Rahul Batra ^78^, Katherine L Bellis ^1, 3^, Beth Blane ^3^, Sophia T Girgis ^3^, Angie Green ^26^, Anita Justice ^28^, Mark Kristiansen ^41^, and Rachel J Williams ^41^.

**Project administration, software and analysis tools:**

Radoslaw Poplawski^15^.

**Project administration and visualisation:**

Garry P Scarlett ^69^.

**Leadership, supervision, and funding acquisition:**

John A Todd ^26^, Christophe Fraser ^27^, Judith Breuer ^40, 41^, Sergi Castellano ^41^, Stephen L Michell ^49^, Dimitris Gramatopoulos ^73^, and Jonathan Edgeworth ^78^.

**Leadership, supervision and metadata curation:**

Gemma L Kay ^51^.

**Leadership, supervision, sequencing and analysis:**

Ana da Silva Filipe ^21^, Aaron R Jeffries ^49^, Sascha Ott ^71^, Oliver Pybus ^24^, David L Robertson ^21^, David A Simpson ^6^, and Chris Williams ^33^.

**Samples, logistics, leadership and supervision:**

Cressida Auckland ^50^, John Boyes ^83^, Samir Dervisevic ^52^, Sian Ellard ^49, 50^, Sonia Goncalves^1^, Emma J Meader ^51^, Peter Muir ^2^, Husam Osman ^95^, Reenesh Prakash ^52^, Venkat Sivaprakasam ^18^, and Ian B Vipond ^2^.

**Leadership, supervision and visualisation**

Jane AH Masoli ^49, 50^.

**Sequencing, analysis and metadata curation**

Nabil-Fareed Alikhan ^51^, Matthew Carlile ^54^, Noel Craine ^33^, Sam T Haldenby ^46^, Nadine Holmes ^54^, Ronan A Lyons ^37^, Christopher Moore ^54^, Malorie Perry ^33^, Ben Warne ^80^, and Thomas Williams ^19^.

**Samples, logistics and metadata curation:**

Lisa Berry ^72^, Andrew Bosworth ^95^, Julianne Rose Brown ^40^, Sharon Campbell ^67^, Anna Casey ^17^, Gemma Clark ^56^, Jennifer Collins ^66^, Alison Cox ^43, 44^, Thomas Davis ^84^, Gary Eltringham ^66^, Cariad Evans ^38, 39^, Clive Graham ^64^, Fenella Halstead ^18^, Kathryn Ann Harris ^40^, Christopher Holmes ^58^, Stephanie Hutchings ^2^, Miren Iturriza-Gomara ^46^, Kate Johnson ^38, 39^, Katie Jones ^72^, Alexander J Keeley ^38^, Bridget A Knight ^49 50^, Cherian Koshy^90^, Steven Liggett ^63^, Hannah Lowe ^81^, Anita O Lucaci ^46^, Jessica Lynch ^25, 29^, Patrick C McClure ^55^, Nathan Moore ^31^, Matilde Mori ^25, 29, 32^, David G Partridge ^38, 39^, Pinglawathee Madona ^43, 44^, Hannah M Pymont ^2^, Paul Anthony Randell ^43, 44^, Mohammad Raza ^38, 39^, Felicity Ryan ^81^, Robert Shaw ^28^, Tim J Sloan ^57^, and Emma Swindells ^65^.

**Sequencing, analysis, Samples and logistics:**

Alexander Adams ^33^, Hibo Asad ^33^, Alec Birchley ^33^, Tony Thomas Brooks ^41^, Giselda Bucca ^93^, Ethan Butcher ^70^, Sarah L Caddy ^13^, Laura G Caller ^2,3, 12^, Yasmin Chaudhry ^11^, Jason Coombes ^33^, Michelle Cronin ^33^, Patricia L Dyal ^41^, Johnathan M Evans ^33^, Laia Fina ^33^, Bree Gatica-Wilcox ^33^, Iliana Georgana ^11^, Lauren Gilbert ^33^, Lee Graham ^33^, Danielle C Groves ^38^, Grant Hall ^11^, Ember Hilvers ^33^, Myra Hosmillo ^11^, Hannah Jones ^33^, Sophie Jones ^33^, Fahad A Khokhar ^13^, Sara Kumziene-Summerhayes ^33^, George MacIntyre-Cockett ^26^, Rocio T Martinez Nunez ^94^, Caoimhe McKerr ^33^, Claire McMurray ^15^, Richard Myers ^7^, Yasmin Nicole Panchbhaya ^41^, Malte L Pinckert ^11^, Amy Plimmer ^33^, Joanne Stockton ^15^, Sarah Taylor ^33^, Alicia Thornton ^7^, Amy Trebes ^26^, Alexander J Trotter ^51^, Helena Jane Tutill ^41^, Charlotte A Williams ^41^, Anna Yakovleva ^11^ and Wen C Yew ^62^.

**Sequencing, analysis and software and analysis tools:**

Mohammad T Alam ^71^, Laura Baxter ^71^, Olivia Boyd ^96^, Fabricia F. Nascimento ^96^, Timothy M Freeman ^38^, Lily Geidelberg ^96^, Joseph Hughes ^21^, David Jorgensen ^96^, Benjamin B Lindsey ^38^, Richard J Orton ^21^, Manon Ragonnet-Cronin ^96^ Joel Southgate ^33, 34,^ and Sreenu Vattipally ^21^.

**Samples, logistics and software and analysis tools:**

Igor Starinskij ^23^.

**Visualisation and software and analysis tools:**

Joshua B Singer ^21^, Khalil Abudahab ^1, 30^, Leonardo de Oliveira Martins ^51^, Thanh Le-Viet ^51^, Mirko Menegazzo ^30^, Ben EW Taylor ^1 30^, and Corin A Yeats ^30^.

**Project Administration:**

Sophie Palmer ^3^, Carol M Churcher ^3^, Alisha Davies ^33^, Elen De Lacy ^33^, Fatima Downing ^33^, Sue Edwards ^33^, Nikki Smith ^38^, Francesc Coll ^97^, Nazreen F Hadjirin ^3^ and Frances Bolt ^44, 45^.

**Leadership and supervision:**

Alex Alderton^1^, Matt Berriman^1^, Ian G Charles ^51^, Nicholas Cortes ^31^, Tanya Curran ^88^, John Danesh^1^, Sahar Eldirdiri ^84^, Ngozi Elumogo ^52^, Andrew Hattersley ^49, 50^, Alison Holmes ^44, 45^, Robin Howe ^33^, Rachel Jones ^33^, Anita Kenyon ^84^, Robert A Kingsley ^51^, Dominic Kwiatkowski ^9^, Cordelia Langford^1^, Jenifer Mason^48^, Alison E Mather ^51^, Lizzie Meadows ^51^, Sian Morgan ^36^, James Price ^44, 45^, Trevor I Robinson ^48^, Giri Shankar ^33^, John Wain ^51^, and Mark A Webber ^51^.

**Metadata curation:**

Declan T Bradley ^5, 6^, Michael R Chapman ^1, 3, 4^, Derrick Crooke ^28^, David Eyre ^28^, Martyn Guest ^34^, Huw Gulliver ^34^, Sarah Hoosdally ^28^, Christine Kitchen ^34^, Ian Merrick ^34^, Siddharth Mookerjee ^44, 45^, Robert Munn ^34^, Timothy Peto ^28^, Will Potter ^52^, Dheeraj K Sethi ^52^, Wendy Smith ^56^, Luke B Snell ^75, 94^, Rachael Stanley ^52^, Claire Stuart ^52^ and Elizabeth Wastenge^20^.

**Sequencing and analysis:**

Erwan Acheson ^6^, Safiah Afifi ^36^, Elias Allara ^2, 3^, Roberto Amato ^1^, Adrienn Angyal ^38^, Elihu Aranday-Cortes ^21^, Cristina Ariani ^1^, Jordan Ashworth ^19^, Stephen Attwood ^24^, Alp Aydin ^51^, David J Baker ^51^, Carlos E Balcazar ^19^, Angela Beckett ^68^ Robert Beer ^36^, Gilberto Betancor ^76^, Emma Betteridge ^1^, David Bibby ^7^, Daniel Bradshaw^7^, Catherine Bresner ^34^, Hannah E Bridgewater ^71^, Alice Broos ^21^, Rebecca Brown ^38^, Paul E Brown ^71^, Kirstyn Brunker ^22^, Stephen N Carmichael ^21^, Jeffrey K. J. Cheng ^71^, Dr Rachel Colquhoun ^19^, Gavin Dabrera ^7^, Johnny Debebe ^54^, Eleanor Drury ^1^, Louis du Plessis ^24^, Richard Eccles ^46^, Nicholas Ellaby ^7^, Audrey Farbos ^49^, Ben Farr ^1^, Jacqueline Findlay ^41^, Chloe L Fisher ^74^, Leysa Marie Forrest ^41^, Sarah Francois ^24^, Lucy R. Frost ^71^, William Fuller^34^, Eileen Gallagher ^7^, Michael D Gallagher ^19^, Matthew Gemmell ^46^, Rachel AJ Gilroy ^51^, Scott Goodwin ^1^, Luke R Green ^38^, Richard Gregory ^46^, Natalie Groves ^7^, James W Harrison ^49^, Hassan Hartman ^7^, Andrew R Hesketh ^93^, Verity Hill ^19^, Jonathan Hubb ^7^, Margaret Hughes^46^, David K Jackson ^1^, Ben Jackson ^19^, Keith James ^1^, Natasha Johnson ^21^, Ian Johnston ^1^, Jon-Paul Keatley ^1^, Moritz Kraemer ^24^, Angie Lackenby ^7^, Mara Lawniczak ^1^, David Lee ^7^, Rich Livett ^1^, Stephanie Lo ^1^, Daniel Mair ^21^, Joshua Maksimovic ^36^, Nikos Manesis ^7^, Robin Manley ^49^, Carmen Manso ^7^, Angela Marchbank ^34^, Inigo Martincorena ^1^, Tamyo Mbisa ^7^, Kathryn McCluggage ^36^, JT McCrone ^19^, Shahjahan Miah ^7^, Michelle L Michelsen ^49^, Mari Morgan ^33^, Gaia Nebbia ^78^, Charlotte Nelson ^46^, Jenna Nichols ^21^, Paola Niola ^41^, Kyriaki Nomikou ^21^, Steve Palmer ^1^, Naomi Park ^1^, Yasmin A Parr ^1^, Paul J Parsons ^38^, Vineet Patel ^7^, Minal Patel ^1^, Clare Pearson ^2, 1^, Steven Platt ^7^, Christoph Puethe ^1^, Mike Quail ^1^, Jayna Raghwani ^24^, Lucille Rainbow ^46^, Shavanthi Rajatileka ^1^, Mary Ramsay ^7^, Paola C Resende Silva ^41, 42^, Steven Rudder ^51^, Chris Ruis ^3^, Christine M Sambles ^49^, Fei Sang ^54^, Ulf Schaefer^7^, Emily Scher ^19^, Carol Scott ^1^, Lesley Shirley ^1^, Adrian W Signell ^76^, John Sillitoe ^1^, Christen Smith ^1^, Dr Katherine L Smollett ^21^, Karla Spellman ^36^, Thomas D Stanton ^19^, David J Studholme ^49^, Grace Taylor-Joyce ^71^, Ana P Tedim ^51^, Thomas Thompson ^6^, Nicholas M Thomson ^51^, Scott Thurston^1^, Lily Tong ^21^, Gerry Tonkin-Hill ^1^, Rachel M Tucker ^38^, Edith E Vamos ^4^, Tetyana Vasylyeva^24^, Joanna Warwick-Dugdale ^49^, Danni Weldon ^1^, Mark Whitehead ^46^, David Williams ^7^, Kathleen A Williamson ^19^, Harry D Wilson ^76^, Trudy Workman ^34^, Muhammad Yasir^51^, Xiaoyu Yu ^19^, and Alex Zarebski ^24^.

**Samples and logistics:**

Evelien M Adriaenssens ^51^, Shazaad S Y Ahmad ^2, 47^, Adela Alcolea-Medina ^59, 77^, John Allan ^60^, Patawee Asamaphan ^21^, Laura Atkinson ^40^, Paul Baker ^63^, Jonathan Ball ^55^, Edward Barton^64^, Mathew A Beale^1^, Charlotte Beaver^1^, Andrew Beggs ^16^, Andrew Bell ^51^, Duncan J Berger ^1^, Louise Berry. ^56^, Claire M Bewshea ^49^, Kelly Bicknell ^70^, Paul Bird ^58^, Chloe Bishop ^7^, Tim Boswell ^56^, Cassie Breen ^48^, Sarah K Buddenborg^1^, Shirelle Burton-Fanning ^66^, Vicki Chalker ^7^, Joseph G Chappell ^55^, Themoula Charalampous ^78, 94^, Claire Cormie^3^, Nick Cortes^29, 25^, Lindsay J Coupland ^52^, Angela Cowell ^48^, Rose K Davidson ^53^, Joana Dias ^3^, Maria Diaz ^51^, Thomas Dibling^1^, Matthew J Dorman^1^, Nichola Duckworth^57^, Scott Elliott^70^, Sarah Essex^63^, Karlie Fallon ^58^, Theresa Feltwell ^8^, Vicki M Fleming ^56^, Sally Forrest ^3^, Luke Foulser^1^, Maria V Garcia-Casado^1^, Artemis Gavriil ^41^, Ryan P George ^47^, Laura Gifford ^33^, Harmeet K Gill ^3^, Jane Greenaway ^65^, Luke Griffith^53^, Ana Victoria Gutierrez^51^, Antony D Hale ^85^, Tanzina Haque ^91^, Katherine L Harper ^85^, Ian Harrison ^7^, Judith Heaney ^89^, Thomas Helmer ^58^, Ellen E Higginson^3^, Richard Hopes ^2^, Hannah C Howson-Wells ^56^, Adam D Hunter ^1^, Robert Impey ^70^, Dianne Irish-Tavares ^91^, David A Jackson^1^, Kathryn A Jackson ^46^, Amelia Joseph ^56^, Leanne Kane ^1^, Sally Kay ^1^, Leanne M Kermack ^3^, Manjinder Khakh ^56^, Stephen P Kidd ^29, 25, 31^, Anastasia Kolyva ^51^, Jack CD Lee ^40^, Laura Letchford ^1^, Nick Levene ^79^, Lisa J Levett ^89^, Michelle M Lister ^56^, Allyson Lloyd ^70^, Joshua Loh ^60^, Louissa R Macfarlane-Smith ^85^, Nicholas W Machin ^2,47^, Mailis Maes ^3^, Samantha McGuigan ^1^, Liz McMinn ^1^, Lamia Mestek-Boukhibar ^41^, Zoltan Molnar ^6^, Lynn Monaghan ^79^, Catrin Moore ^27^, Plamena Naydenova ^3^, Alexandra S Neaverson ^1^, Rachel Nelson ^1^, Marc O Niebel ^21^, Elaine O’Toole^48^, Debra Padgett ^64^, Gaurang Patel ^1^, Brendan AI Payne ^66^, Liam Prestwood ^1^, Veena Raviprakash ^67^, Nicola Reynolds^86^, Alex Richter ^16^, Esther Robinson ^95^, Hazel A Rogers^1^, Aileen Rowan ^96^, Garren Scott ^64^, Divya Shah ^40^, Nicola Sheriff ^67^, Graciela Sluga, Emily Souster^1^, Michael Spencer-Chapman^1^, Sushmita Sridhar ^1, 3^, Tracey Swingler ^53^, Julian Tang^58^, Graham P Taylor^96^, Theocharis Tsoleridis ^55^, Lance Turtle^46^, Sarah Walsh ^57^, Michelle Wantoch ^86^, Joanne Watts ^48^, Sheila Waugh ^66^, Sam Weeks^41^, Rebecca Williams^31^, Iona Willingham^56^, Emma L Wise ^25, 29, 31^, Victoria Wright ^54^, Sarah Wyllie ^70^, and Jamie Young ^3^.

**Software and analysis tools**

Amy Gaskin^33^, Will Rowe ^15^, and Igor Siveroni ^96^.

**Visualisation:**

Robert Johnson ^96^.

**1** Wellcome Sanger Institute, **2** Public Health England, **3** University of Cambridge, **4** Health Data Research UK, Cambridge, **5** Public Health Agency, Northern Ireland, **6** Queen’s University Belfast **7** Public Health England Colindale, **8** Department of Medicine, University of Cambridge, **9** University of Oxford, **10** Departments of Infectious Diseases and Microbiology, Cambridge University Hospitals NHS Foundation Trust; Cambridge, UK, **II** Division of Virology, Department of Pathology, University of Cambridge, **12** The Francis Crick Institute, **13** Cambridge Institute for Therapeutic Immunology and Infectious Disease, Department of Medicine, **14** Public Health England, Clinical Microbiology and Public Health Laboratory, Cambridge, UK, **15** Institute of Microbiology and Infection, University of Birmingham, **16** University of Birmingham, **17** Queen Elizabeth Hospital, **18** Heartlands Hospital, **19** University of Edinburgh, **20** NHS Lothian, **21** MRC-University of Glasgow Centre for Virus Research, **22** Institute of Biodiversity, Animal Health & Comparative Medicine, University of Glasgow, **23** West of Scotland Specialist Virology Centre, **24** Dept Zoology, University of Oxford, **25** University of Surrey, **26** Wellcome Centre for Human Genetics, Nuffield Department of Medicine, University of Oxford, **27** Big Data Institute, Nuffield Department of Medicine, University of Oxford, **28** Oxford University Hospitals NHS Foundation Trust, **29** Basingstoke Hospital, **30** Centre for Genomic Pathogen Surveillance, University of Oxford, **31** Hampshire Hospitals NHS Foundation Trust, **32** University of Southampton, **33** Public Health Wales NHS Trust, **34** Cardiff University, **35** Betsi Cadwaladr University Health Board, **36** Cardiff and Vale University Health Board, **37** Swansea University, **38** University of Sheffield, **39** Sheffield Teaching Hospitals, **40** Great Ormond Street NHS Foundation Trust, **41** University College London, **42** Oswaldo Cruz Institute, Rio de Janeiro **43** North West London Pathology, **44** Imperial College Healthcare NHS Trust, **45** NIHR Health Protection Research Unit in HCAI and AMR, Imperial College London, **46** University of Liverpool, **47** Manchester University NHS Foundation Trust, **48** Liverpool Clinical Laboratories, **49** University of Exeter, **50** Royal Devon and Exeter NHS Foundation Trust, **51** Quadram Institute Bioscience, University of East Anglia, **52** Norfolk and Norwich University Hospital, **53** University of East Anglia, **54** Deep Seq, School of Life Sciences, Queens Medical Centre, University of Nottingham, **55** Virology, School of Life Sciences, Queens Medical Centre, University of Nottingham, **56** Clinical Microbiology Department, Queens Medical Centre, **57** PathLinks, Northern Lincolnshire & Goole NHS Foundation Trust, **58** Clinical Microbiology, University Hospitals of Leicester NHS Trust, **59** Viapath, **60** Hub for Biotechnology in the Built Environment, Northumbria University, **61** NU-OMICS Northumbria University, **62** Northumbria University, **63** South Tees Hospitals NHS Foundation Trust, **64** North Cumbria Integrated Care NHS Foundation Trust, **65** North Tees and Hartlepool NHS Foundation Trust, **66** Newcastle Hospitals NHS Foundation Trust, **67** County Durham and Darlington NHS Foundation Trust, **68** Centre for Enzyme Innovation, University of Portsmouth, **69** School of Biological Sciences, University of Portsmouth, **70** Portsmouth Hospitals NHS Trust, **71** University of Warwick, **72** University Hospitals Coventry and Warwickshire, **73** Warwick Medical School and Institute of Precision Diagnostics, Pathology, UHCW NHS Trust, **74** Genomics Innovation Unit, Guy’s and St. Thomas’ NHS Foundation Trust, **75** Centre for Clinical Infection & Diagnostics Research, St. Thomas’ Hospital and Kings College London, **76** Department of Infectious Diseases, King’s College London, **77** Guy’s and St. Thomas’ Hospitals NHS Foundation Trust, **78** Centre for Clinical Infection and Diagnostics Research, Department of Infectious Diseases, Guy’s and St Thomas’ NHS Foundation Trust, **79** Princess Alexandra Hospital Microbiology Dept., **80** Cambridge University Hospitals NHS Foundation Trust, **81** East Kent Hospitals University NHS Foundation Trust, **82** University of Kent, **83** Gloucestershire Hospitals NHS Foundation Trust, **84** Department of Microbiology, Kettering General Hospital, **85** National Infection Service, PHE and Leeds Teaching Hospitals Trust, **86** Cambridge Stem Cell Institute, University of Cambridge, **87** Public Health Scotland, 88 Belfast Health & Social Care Trust, **89** Health Services Laboratories, **90** Barking, Havering and Redbridge University Hospitals NHS Trust, **91** Royal Free NHS Trust, **92** Maidstone and Tunbridge Wells NHS Trust, **93** University of Brighton, **94** Kings College London, **95** PHE Heartlands, **96** Imperial College London, **97** Department of Infection Biology, London School of Hygiene and Tropical Medicine.

